# Spatiotemporal manipulation of the mismatch repair system of *Pseudomonas putida* accelerates phenotype emergence

**DOI:** 10.1101/2021.01.21.427673

**Authors:** Lorena Fernández-Cabezón, Antonin Cros, Pablo I. Nikel

## Abstract

Developing complex phenotypes in industrially-relevant bacteria is a major goal of metabolic engineering, which encompasses the implementation of both rational and random approaches. In the latter case, several tools have been developed towards increasing mutation frequencies—yet the precise spatiotemporal control of mutagenesis processes continues to represent a significant technical challenge. *Pseudomonas* species are endowed with one of the most efficient DNA mismatch repair (MMR) systems found in bacteria. Here, we investigated if the endogenous MMR system could be manipulated as a general strategy to artificially alter mutation rates in *Pseudomonas* species. To bestow a conditional mutator phenotype in the platform bacterium *Pseudomonas putida*, we constructed inducible mutator devices to modulate the expression of the dominant-negative *mutL*^*E36K*^ allele. Regulatable overexpression of *mutL*^*E36K*^ in a broad-host-range, easy-to-cure plasmid format resulted in a transitory inhibition of the MMR machinery, leading to a significant increase (up to 438-fold) in mutation frequencies and a heritable fixation of genome mutations. Following such accelerated mutagenesis-followed-by selection approach, three phenotypes were successfully evolved: resistance to antibiotics streptomycin and rifampicin and reversion of a synthetic uracil auxotrophy. Thus, these mutator devices could be applied to accelerate evolution of metabolic pathways in long-term evolutionary experiments, alternating cycles of (inducible) mutagenesis coupled to selection schemes.

## INTRODUCTION

Systems metabolic engineering and synthetic biology guide the development of microbial cell factories (MCFs) capable of converting renewable raw materials into value-added compounds^1-4^. However, low productivities and product yields by most MCFs, even after comprehensive optimization of biosynthetic pathways, continue to make the implementation of economically-viable bioprocesses difficult at an industrial scale^5^. Low product yields are often caused by a decrease in cell viability and genetic instability of MCFs under industrially-relevant production conditions^6-7^. For instance, the presence of growth inhibitors in renewable raw materials (e.g. crude glycerol and biomass hydrolysates) and the accumulation of toxic compounds during fermentation (including metabolic intermediates and target products) are known to negatively impact cell survival^8-10^.

Adaptive laboratory evolution (ALE), also known as *evolutionary engineering*, is a valuable tool to improve complex phenotypic traits that can be coupled with microbial growth (e.g. tolerance to inhibitors, substrate utilization, growth temperature)^11-13^. At its core, ALE involves the extended propagation of a microbial strain or population, typically for hundreds of generations, in the presence of a desired selective pressure. Mutants that accumulate beneficial mutations will occasionally emerge and expand within the population over time. Selected mutants displaying enhanced phenotypes can be subsequently characterized and sequenced towards reverse engineering^11-16^. Unlike purely rational approaches, ALE facilitates the identification of non-intuitive beneficial mutations that occur in a variety of genes in parallel without requiring any knowledge of underlying genetic mechanisms.

Since intrinsic DNA mutation rates are typically very low (ranging in the order of 10^−9^–10^−10^ per base pair per generation)^17-18^, small and transient increases in mutation frequency can significantly improve the accumulation of beneficial mutations in microbial populations^13,19^. This rationale has been applied to certain ALE experiments in which the genetic diversity of a microbial population was increased before and/or during growth under restrictive culture conditions^6,16,20-21^. Chemical and/or physical mutagenesis techniques have been traditionally used due to their simplicity and wide applicability^22-23^, but other genome-wide random mutagenesis techniques can be also applied for this purpose^6^. Mutator strains, i.e. bacteria displaying higher mutation rates, frequently have mutations in one or several genes encoding DNA repair or error-avoidance systems^24^. Most bacteria control DNA substitution rates through overlapping DNA repair mechanisms, subdivided into three main categories: (i) base selection, (ii) proofreading and (iii) mismatch repair (MMR)^25^. Base selection encompasses the discrimination between correct and incorrect nucleotides by DNA polymerase, while proofreading is the subsequent editing of the newly incorporated nucleotide by a 3’→5’ exonuclease activity that hydrolyzes incorrect bases. Following replication, newly replicated DNA is checked by a MMR system that recognizes and corrects mismatches resulting from replication errors^25^. Specific mutations in components of MMR (e.g. *mutL* and *mutS*) or in proofreading DNA polymerases (e.g. *dnaQ*), as well as the overexpression of certain dominant-negative mutator alleles of the same genes, have been shown to result in mutator phenotypes^24,26^. Conditional mutator phenotypes have been applied to the phenotypic optimization of MCFs over time^27-32^. However, a major problem of these conditional phenotypes is the relatively low control of spatiotemporal activity afforded by the cognate devices. A typical problem of these systems is that the ability of effectively halting mutagenesis is limited, and the cells will continue to mutate even after a desired phenotype is achieved^33^.

*Pseudomonas putida* is a ubiquitous Gram-negative bacterium used for biotechnological and bioremediation applications^34-37^. Strain KT2440, for instance, is a promising microbial *chassis* for handling the synthesis of difficult-to-produce chemicals involving harsh reactions and complex biochemistries^36-40^. Alas, metabolic engineering of *P. putida* still relies largely on trial-and-error approaches. While advanced genome-wide engineering tools are being constantly developed and optimized^41-44^, complex phenotypes are the result of multi-level regulatory layers that are often difficult to design from first principles. ALE has recently started to be exploited in *P. putida*-based MCFs^45-50^. On this background, we set out to explore if genome-wide mutation rates in *P. putida* (both wild-type strain and reduced-genome derivatives thereof) could be increased by synthetic control of the well- characterized MMR in this bacterium^51^. To this end, in this work we have designed a toolbox to conditionally increase mutation rates in Gram-negative bacteria by specifically interfering with the endogenous MMR system towards accelerating the evolution of specific phenotypes. Moreover, we focused on the adoption of emerging strategies to easily cure plasmid-born mutator devices from bacterial populations, such that the temporal window of increased mutagenesis rates can be externally controlled. The application of this set of synthetic mutator devices has been systematically validated in evolution experiments targeting both antibiotic resistance and growth phenotypes *via* auxotrophy reversion.

## RESULTS AND DISCUSSION

### Construction of broad-host-range, plasmid-based mutator devices to increase DNA mutation rates

*Pseudomonas* species have been shown to display one of the highest MMR efficiencies found in bacteria (e.g. *P. fluorescens*^52^). Therefore, we hypothesized that manipulating the endogenous MMR system could be a straightforward approach to increase the mutation rate in bacterial species of the *Pseudomonas* genus. In order to bestow a conditional mutator phenotype in our model bacterium *P. putida*, we constructed two inducible mutator devices, based on well-characterized expression systems, to tightly modulate the expression of the mutator allele *mutL*^*E36K*^ from *P. putida*^51^ (**Fig. 1** and **Fig. S1** in the Supporting Information). The E36K amino acid change in MutL stems from a 106(G→A) mutation in the corresponding allele (**Fig. S1**). The overexpression of the homologous, dominant-negative allele *mutL*^*E32K*^ from *Escherichia coli* has been shown to result in a transitory inhibition of the MMR machinery^26,53^, which leads to the heritable fixation of mutations in the genome by tampering with the MMR system (**Fig. 1a**). The *mutL*^*E36K*^ allele, in contrast, has been exploited for genome engineering approaches specifically developed for *P. putida* and related species^54^. In our mutator devices, the expression of *mutL*^*E36K*^ was driven from two tightly-regulated expression systems, i.e. the thermoinducible *c*I857/P_L_ expression system from the bacteriophage λ and the cyclohexanone-inducible ChnR/P_*chnB*_ system from *Acinetobacter johnsonii*. Both expression vectors have been previously employed for heterologous gene expression in Gram-negative bacteria such as *E. coli* or *P. putida*^55-58^. Thus, the *mutL*^*E36K*^ gene was cloned into vectors pSEVA2514 (*c*I857/P_L_) and pSEVA2311 (ChnR/P_*chnB*_) to yield the mutator plasmids pS2514M and pS2311M, respectively (**Fig. 1b**). By adopting the rules set in the *Standard European Vector Architecture* (SEVA) platform^59^, the subsequent transfer of the mutator devices and plasmids developed herein to various bacterial hosts is greatly facilitated. Moreover, the implementation of these two expression systems enables the user to decide whether induction of the system can be done by a temperature shift (to 40°C) or addition of chemicals to the culture medium (cyclohexanone). These two approaches were selected as the first one (*c*I857/P_L_) relies on relieving the transcriptional repression mediated by the *c*I857 protein when it gets degraded at 40°C, whereas the ChnR/P_*chnB*_ system acts *via* direct activation of the transcriptional response upon addition of the small-molecule inducer (**Fig. 1c**).

**Figure 1.**
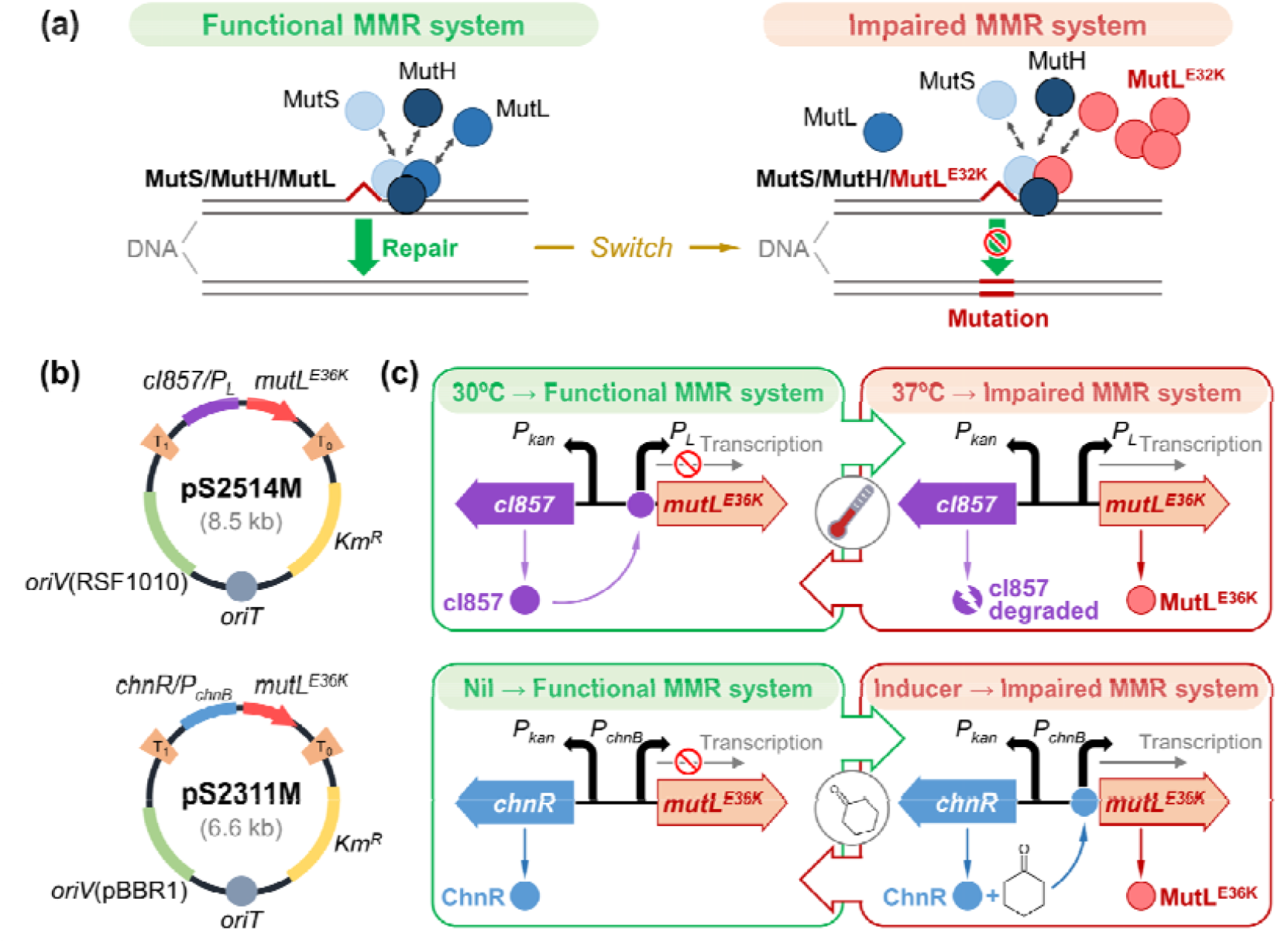
Construction of broad-host-range mutator devices to conditionally increasing mutation rates in Gram-negative bacteria. (a) The bacterial DNA mismatch repair (MMR) system recognizes and fixes mutations that arise during DNA replication and recombination. MutS recognizes genomic DNA mismatches and recruits MutL. The MutL/MutS complex activates the MutH endonuclease, which cleaves the newly synthesized, unmethylated daughter strand at the nearest hemimethylated d(GATC) site, and thereby marks it for a removal and a repair–synthesis process that involves a variety of other proteins. Overexpression of the dominant-negative mutator allele *mutL*^*E32K*^ from *E. coli* increases mutation rates^26^. **(b)** Structure of the two mutator devices used in this work. Plasmids pS2514M and pS2311M, based on the *Standard European Vector Architecture*^90^, were designed for thermo-inducible or cyclohexanone-inducible expression of the mutator allele *mutL*^*E36K*^ from *P. putida*, respectively. Functional elements in the plasmids not drawn to scale; *Km*^R^, kanamycin- resistance marker. **(c)** Two strategies for tampering with the MMR system of *P. putida*. When using plasmid pS2514M, the temperature-sensitive repressor *c*I857 is constitutively produced at 28-32°C and specifically binds to the *P*_*L*_ promoter, mediating transcriptional repression of the gene cloned downstream (i.e. *mutL*^*E36K*^). By shifting the temperature above 37°C (e.g. 40°C), the expression of *mutL*^*E36K*^ takes place due to the denaturation of *c*I857. When using plasmid pS2311M, the ChnR transcriptional regulator is constitutively synthetized and binds to the *P*_*chnB*_ promoter in the presence of its inducer (cyclohexanone), thus causing the expression of the gene cloned downstream (i.e. *mutL*^*E36K*^).

### Emergence of antibiotic resistance phenotypes in *P. putida* carrying synthetic mutator devices

To investigate the functionality of the mutator devices, the occurrence of antibiotic-resistant mutants was assessed in bacterial cultures grown in liquid medium. Two types of antibiotic resistance were selected to this end, namely, rifampicin (Rif) and streptomycin (Str), and the systems were firstly calibrated with the wild-type strain KT2440. In these experiments, control strains (i.e. *P. putida* KT2440/pSEVA2514 and KT2440/pSEVA2311) and their derivatives carrying the conditional mutator devices (i.e. *P. putida* KT2440/pS2514M and KT2440/pS2311M) were cultured at 30°C in non-selective M9 minimal medium containing glucose, and subjected to a mutagenesis protocol as indicated in **Fig. 2** and *Methods*. In the case of strains carrying vectors with the *c*I857/P_L_ expression system, cultures were shifted at 40°C for 15 min for induction; whereas cyclohexanone was added at 1 mM in cultures of the strains transformed with vectors bearing the ChnR/P_*chnB*_ system. Cultures were re-incubated at 30 °C, after temporally inducing a mutator phenotype, and were stopped at different phases of bacterial growth (i.e. early-exponential, mid-exponential or stationary phase) to assess the appearance of the target phenotypes.

**Figure 2.**
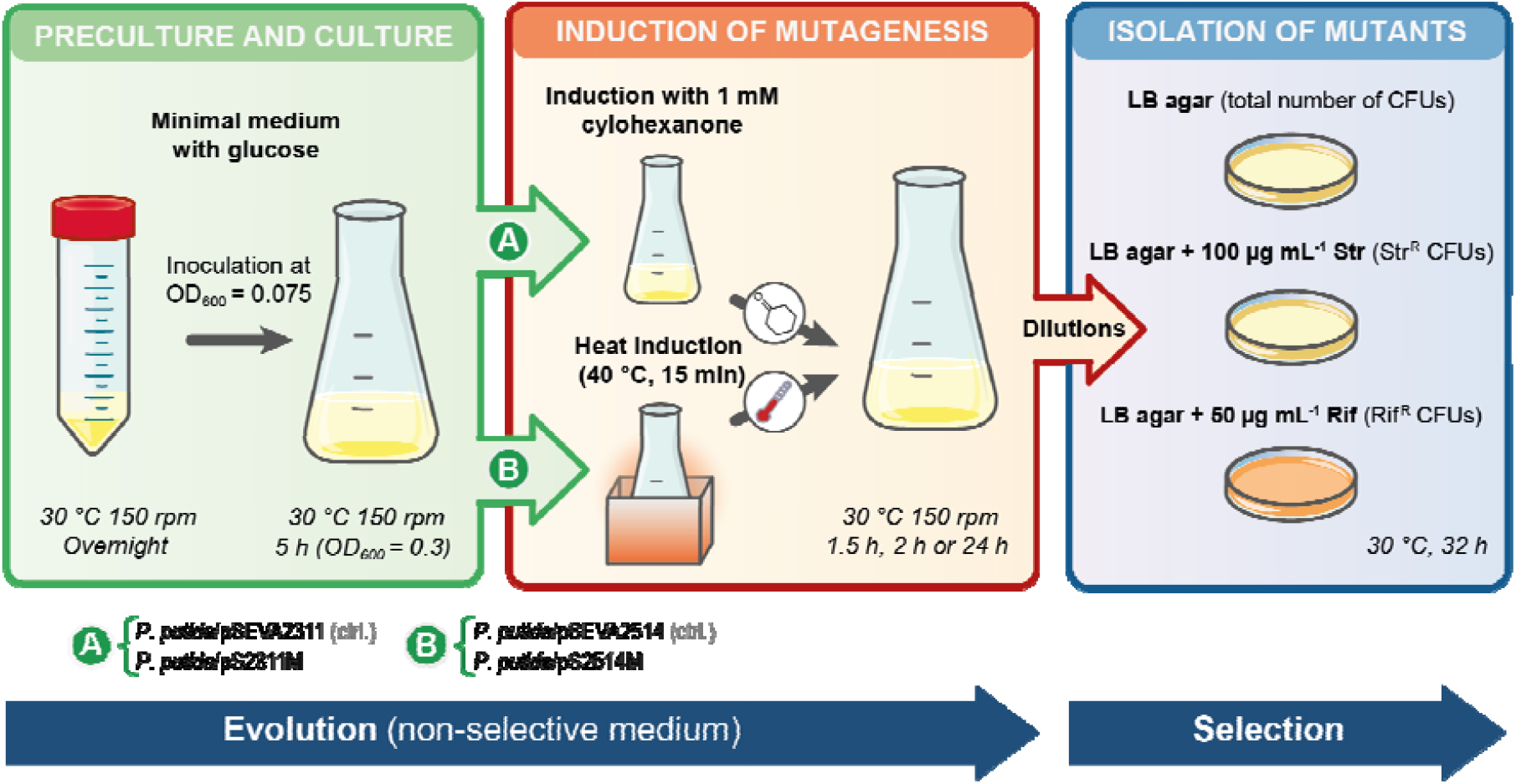
Experimental setup for evolution of P. putida strains carrying mutator plasmids. Control (ctrl.) strains (i.e. *P. putida* KT2440/pSEVA2514 and KT2440/pSEVA2311, carrying empty vectors) and conditional-mutator strains (i.e. *P. putida* KT2440/pS2514M and KT2440/pS2311M) were incubated in shaken-flask cultures in a non-selective medium [e.g. M9 minimal medium containing 0.3% (w/v) glucose]. After 5 h, when cultures reached an optical density at 600 nm (OD_600_) = 0.3, the expre*s*sion systems were induced either thermally (incubating the flasks in a water bath) or chemically (adding cyclohexanone to the medium). The cultures were re-incubated at 30°C with shaking and stopped after 1.5 h (early exponential phase, OD_600_ = 0.6), 2 h (mid-exponential phase, OD_600_ = 1) or 24 h (stationary phase). Several aliquots cultures were plated onto a selective solid medium [e.g. LB agar supplemented with streptomycin (Str) or rifampicin (Rif)] to assess the appearance of mutants in the bacterial population [e.g. rifampicin- (Rif^R^) or streptomycin-resistant (Str^R^) mutants]. The total number of viable cells in the bacterial cultures was estimated by plating dilutions of the cultures on non-selective solid medium (e.g. LB agar). *CFU*, colony-forming unit.

The occurrence of mutants developing resistance to either Rif or Str was investigated in the bacterial populations after the treatments indicated above. Resistance to these antibiotics has been widely used for the investigation of spontaneous and induced mutagenesis processes in Gram-negative bacteria^60-62^. Rifampin-resistant (Rif^R^) and streptomycin-resistant (Str^R^) phenotypes occur due to the appearance of mutations in the *rpoB* and *rpsL* genes, encoding the β-subunit of RNA polymerase and the 30S ribosomal protein S12, respectively^60-63^. Mutation frequencies were estimated by assessing the frequency of occurrence of Rif^R^ or Str^R^ cells on the total number of viable cells in the bacterial population for each tested experimental condition (**Fig. 3**). In all accelerated mutagenesis experiments, we observed a significantly higher number of Rif^R^ and Str^R^ mutants isolated in selective conditions in bacterial clones carrying a mutator device compared to their respective control strains (**Fig. 3**). A visual example of this general trend is presented in **Fig. S2** in the Supporting Information. The number of Rif^R^ and Str^R^ colonies present in 5 mL of non-diluted cultures of *P. putida* KT2440/pSEVA2311 (plated after concentrating the biomass by centrifugation and resuspension) was roughly similar to that in selective plates seeded with only 100 μL of an undiluted culture of *P. putida* KT2440/pS2311M. When these differences were properly quantified, we observed that the frequency of appearance of Rif^R^ and Str^R^ mutants in *P. putida* KT2440/pS2311M was 438- and 10-fold higher, respectively, as compared to the control strain when the induction of the expression system was stopped in early-exponential growth phase (**Fig. 3a**). In the same experimental conditions, the frequency of occurrence of Rif^R^ and Str^R^ mutants in *P. putida* KT2440/pS2514M was 45- and 14-fold higher compared to the control strain, respectively (**Fig. 3b**). Similar relative mutation frequencies were observed when the cultures of the different recombinant strains were prolonged until reaching mid-exponential and stationary phase (**Fig. 3**). The largest differences in mutation frequencies were observed in actively-growing cells (i.e. during the early- or mid-exponential phase of growth) as compared to bacteria harvested during the stationary phase^64-66^.

**Figure 3.**
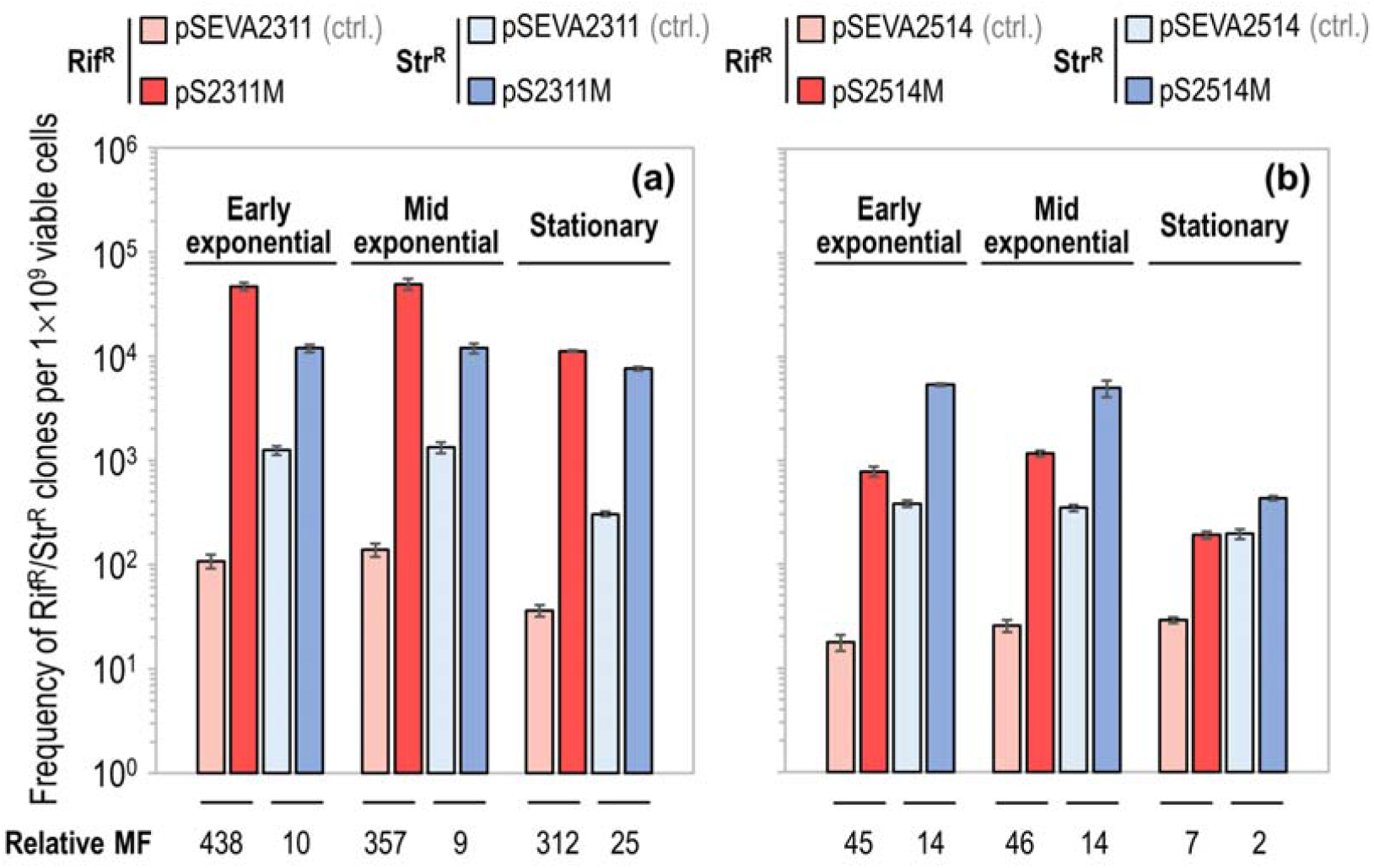
Evolution of antibiotic resistance in *P. putida* using conditional mutator devices. *P. putida* strains carrying the systems inducible by cyclohexanone **(a)** and temperature shifts **(b)** were evolved by following the mutagenesis protocol described in **Fig. 2**. Culture aliquots were plated onto selective medium [i.e. LB agar supplemented with 100 μg mL^−1^ streptomycin (Str) or 50 μg mL^−1^ rifampicin (Rif)] to determine the appearance of Rif- (Rif^R^) or Str-resistant (Str^R^) mutants in the population after evolution. The total number of viable cells was estimated by plating dilutions of each of the cultures onto LB agar plates. Two technical replicates and several dilutions for replicate were performed for each bacterial strain and per each selective culture condition. Columns represent mean values of mutation frequencies (MF, expressed as the number of mutant cells per 10^9^ viable *P. putida* cells) from at least two independent experiments ± standard deviation. *Relative mutation frequencies* were obtained by comparing the mutation frequency of the conditional mutator strain with the respective control (ctrl.) strain in the same experimental setup (i.e. expressed as fold-change).

Taken together, these experimental data demonstrate the functionality of the mutator devices developed in this work to temporarily increase the global mutation rate in *P. putida*. The differences detected in the mutation frequencies as elicited by the two mutator devices may be related to intrinsic properties of each of the plasmids that carry the mutator allele (e.g. origin of replication and promoter used, since this will affect the transcriptional output), as well as to the protocols followed to induce the expression of *mutL*^*E36K*^. Moreover, differences in mutation frequencies are known to arise depending on the method used for their estimation (i.e. counting the occurrence of Rif^R^ or Str^R^ clones). On the one hand, mutation frequencies and the actual spectrum of mutations have been shown to vary at different chromosomal positions in several bacterial species, including *P. putida*^67-69^. Other genetic factors, such as the orientation of the target gene in the replication fork, its level of transcription and/or the immediately flanking nucleotides can also influence the mutation frequency^67-68^. On the other hand, the nature of the mutations acquired by *rpoB* and *rpsL* has been demonstrated to lead to distinct levels of resistance to both Rif and Str, which makes it difficult to use these phenotypes for a direct, quantitative estimation of global mutation rates in different bacterial strains. Factors such as the time and temperature of incubation in selective medium (i.e. agar plates supplemented with antibiotic) have been also shown to dramatically affect the estimation of mutation frequencies (e.g. due to the appearance of colonies with uneven sizes)^62^. Therefore, the utilization of alternative phenotypes is highly recommendable for the calibration and validation of our mutator tool. This issue was undertaken as explained in the next section.

### Reversion of a uracil auxotrophy in *P. putida* using mutator devices

To further calibrate the mutator vectors and gain insight into growth phenotypes beyond antibiotic resistance, we investigated the reversion of uracil auxotrophy of the *P. putida pyrF* HM (**Table 1**). This strain is a derivative of reduced-genome *P. putida* EM42 carrying a 163(A→T) mutation in *pyrF*, which results in a Lys55*STOP* change in the PyrF protein^70^. This change, in turn, leads to abortive translation of the cognate mRNA and the strain thus lacks a functional orotidine 5’-phosphate decarboxylase (i.e. Ura^−^ phenotype), an essential activity for bacterial growth on minimal medium. In these experiments, the control strains, i.e. *P. putida pyrF* HM/pSEVA2514 and *pyrF* HM/pSEVA2311, and the conditional mutator strains, i.e. *P. putida pyrF* HM/pS2514M and *pyrF* HM/pS2311M, were cultured at 30°C in non-selective medium (i.e. with uracil supplementation) and subjected to the mutagenesis protocol indicated in **Fig. 2**. After treatment, the cultures were re-incubated at 30 °C, and were harvested upon a doubling in the population size (i.e. early-exponential phase). The emergence of uracil prototrophic mutants (Ura^+^) in the evolved bacterial populations was determined by seeding M9 minimal medium agar plates with glucose but without uracil supplementation. Mutation frequencies were estimated by assessing the frequency of occurrence of Ura^+^ mutants on the total number of viable cells in the population for each tested experimental condition (**Fig. 4a**). A significant higher number of Ura^+^ mutants were isolated from the bacterial populations carrying the conditional mutator devices as compared to their respective control strains, again validating the functionality of the mutator tools. In fact, we only isolated a negligible number (0-4) of spontaneous Ura^+^ mutants in bacterial populations of control strains under these experimental conditions. Under these conditions, the devices borne by the mutator plasmids pS2311M and pS2514M mediated an increase in the relative mutation frequency of 51- and 384-fold, respectively, as compared to control conditions. Interestingly, in this case no significant differences were found when comparing mutation frequencies estimated for the cyclohexanone inducible and thermoinducible mutator systems (i.e. 750 and 860 Ura^+^ mutants per 10^9^ viable cells, respectively, **Fig. 4a**). The next objective in this experiment was studying the nature of the mutations acquired by the Ura^+^ clones.

**Table 1.**
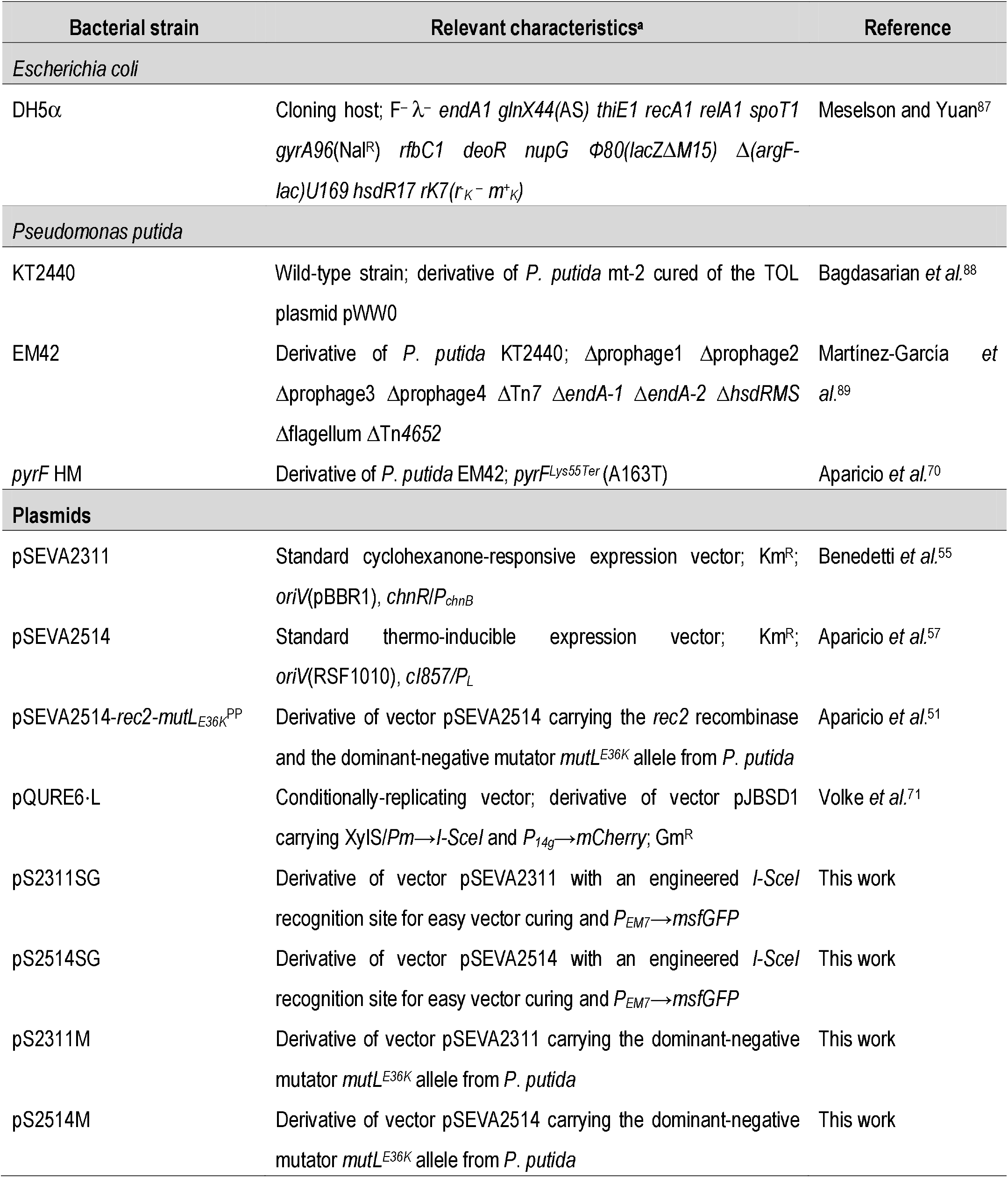
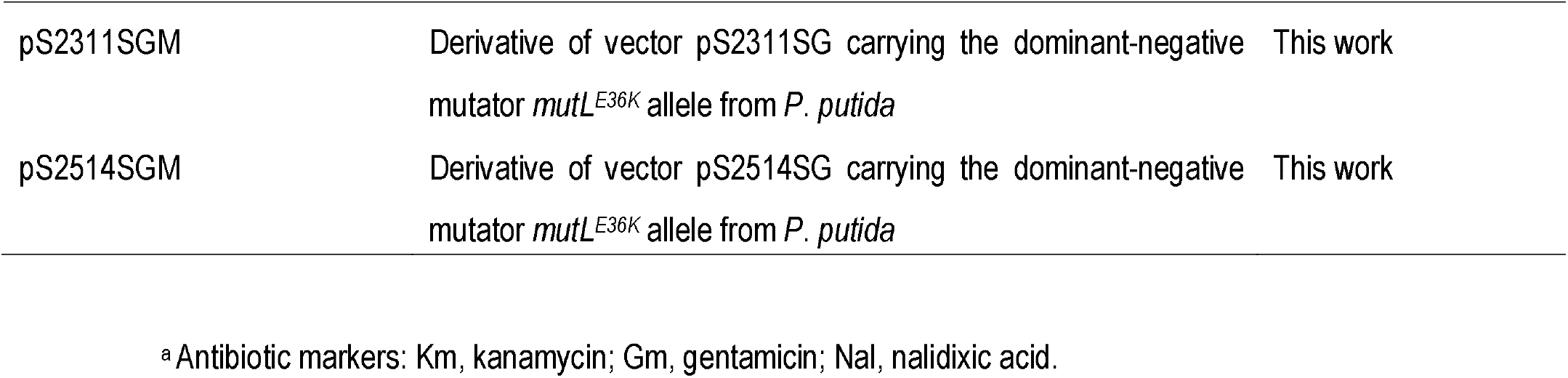
Bacterial strains and plasmids used in this work.

| Bacterial strain | Relevant characteristics<sup>a</sup> | Reference |
| --- | --- | --- |
| *Escherichia coli* |  |  |
| DH5α | Cloning host; F-λ-*endA1 glnX44(AS) thiE1 recA1 relA1 spoT1 gyrA96(Nal<sup>R</sup>) rfbC1 deoR nupG Φ80(lacZΔM15) $\Delta$(argF-lac)U169 hsdR17 rK7(r<sub>K</sub>-m<sup>+</sup><sub>K</sub>)* | Meselson and Yuan<sup>87</sup> |
| *Pseudomonas putida* |  |  |
| KT2440 | Wild-type strain; derivative of *P. putida* mt-2 cured of the TOL plasmid pWW0 | Bagdasarian *et al.*<sup>88</sup> |
| EM42 | Derivative of *P. putida* KT2440; Δprophage1 Δprophage2 Δprophage3 Δprophage4 ΔTn7 ΔendA-1 ΔendA-2 ΔhsdRMS Δflagellum ΔTn4652 | Martínez-García *et al.*<sup>89</sup> |
| *pyrF* HM | Derivative of *P. putida* EM42; *pyrF*<sup>Lys55Ter</sup> (A163T) | Aparicio *et al.*<sup>70</sup> |
| **Plasmids** |  |  |
| pSEVA2311 | Standard cyclohexanone-responsive expression vector; Km<sup>R</sup>; *oriV*(pBBR1), *chnR/P<sub>chnB</sub>* | Benedetti *et al.*<sup>55</sup> |
| pSEVA2514 | Standard thermo-inducible expression vector; Km<sup>R</sup>; *oriV*(RSF1010), *cl857/P<sub>L</sub>* | Aparicio *et al.*<sup>57</sup> |
| pSEVA2514-*rec2-mutL*<sup>E36K</sup>PP | Derivative of vector pSEVA2514 carrying the *rec2* recombinase and the dominant-negative mutator *mutL*<sup>E36K</sup> allele from *P. putida* | Aparicio *et al.*<sup>51</sup> |
| pQURE6-L | Conditionally-replicating vector; derivative of vector pJBSD1 carrying *XylS/Pm*→*I-SceI* and *P<sub>14g</sub>*→*mCherry*; Gm<sup>R</sup> | Volke *et al.*<sup>71</sup> |
| pS2311SG | Derivative of vector pSEVA2311 with an engineered *I-SceI* recognition site for easy vector curing and *P<sub>EM7</sub>*→*msfGFP* | This work |
| pS2514SG | Derivative of vector pSEVA2514 with an engineered *I-SceI* recognition site for easy vector curing and *P<sub>EM7</sub>*→*msfGFP* | This work |
| pS2311M | Derivative of vector pSEVA2311 carrying the dominant-negative mutator *mutL*<sup>E36K</sup> allele from *P. putida* | This work |
| pS2514M | Derivative of vector pSEVA2514 carrying the dominant-negative mutator *mutL*<sup>E36K</sup> allele from *P. putida* | This work |
| pS2311SGM | Derivative of vector pS2311SG carrying the dominant-negative mutator *mutL*<sup>E36K</sup> allele from *P. putida* | This work |
| pS2514SGM | Derivative of vector pS2514SG carrying the dominant-negative mutator *mutL*<sup>E36K</sup> allele from *P. putida* | This work |

<sup>a</sup> Antibiotic markers: Km, kanamycin; Gm, gentamicin; Nal, nalidixic acid.

**Figure 4.**
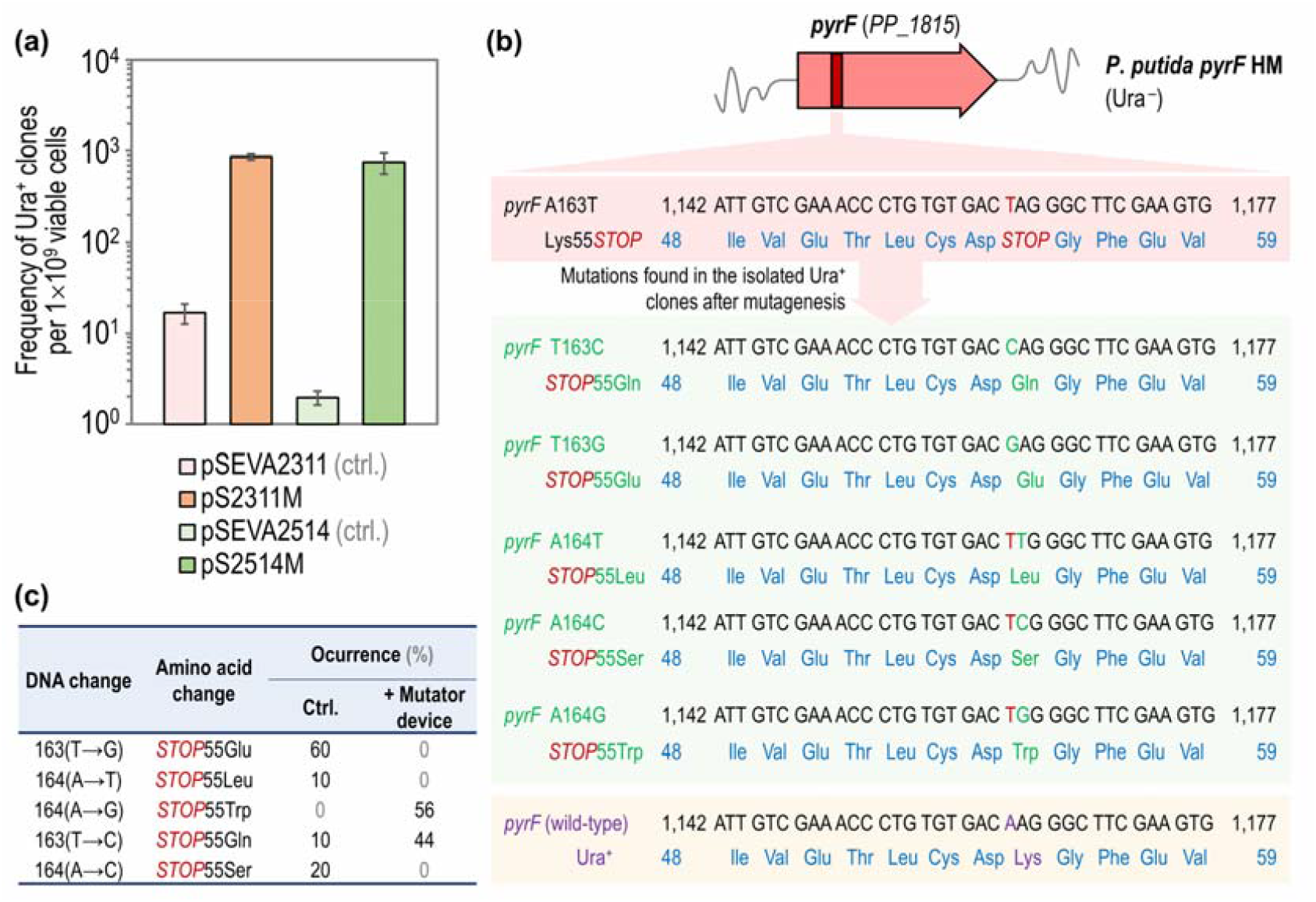
Reversion of the uracil auxotrophy in *P. putida pyrF* using mutator devices. (a)*P. putida pyrF* HM, carrying the *pyrF*^Lys55STOP^ allele that confers uracil auxotrophy (Ura^−^), was transformed with the two conditional mutator systems (i.e. inducible by cyclohexanone or temperature shifts) or the corresponding control (ctrl.) vectors, and evolved by following the mutagenesis protocol described in **Fig. 2**. Several aliquots of these bacterial cultures were plated on selective solid medium (i.e. M9 minimal medium containing glucose as the only carbon source) to estimate the appearance of uracil prototrophic mutants (Ura^+^) in the population. The total number of viable cells was estimated by plating dilutions of the same cultures onto M9 minimal medium plates supplemented with glucose and 20 μg mL^−1^ uracil. Two technical replicates and several dilutions for replicate were performed for each bacterial strain and per each selective culture condition. Columns represent mean values of mutation frequencies (expressed as the number of mutant cells per 10^9^ viable *P. putida pyrF* HM cells) from at least two independent experiments ± standard deviation. **(b)** Mutations found in the *pyrF* gene (*PP_1815*) in the isolated Ura^+^ mutants. **(c)** Frequency of mutation occurrence in control (ctrl.) and in the strain carrying the conditional mutator devices. Stop codons are indicated with the abbreviation *STOP*.

### The conditional mutator phenotype favors the emergence of transition mutations in the genome

To investigate the nature of the mutations introduced with the mutator devices, the whole *pyrF* gene (*PP_1815*) was amplified by high-fidelity PCR from several Ura^+^ clones and the resulting amplicons were sent for sequencing (**Fig. 4b**). Firstly, we isolated multiple Ura^+^ mutants from two independent evolution experiments performed with the conditional mutator strains (i.e. *P. putida pyrF* HM/pS2514M and *pyrF* HM/pS2311M). The DNA transitions 164(A→G) or 163(T→C), which eliminate the premature *STOP* codon in the *pyrF*^163(A→T)^ variant that leads to uracil auxotrophy, were equally frequent in the mutants analyzed across different independent experiments (**Fig. 4b** and **4c**). These observations indicate that the mutator devices seem to mediate changes from a purine nucleotide to another purine (i.e. A:T ↔ G:C) or a pyrimidine nucleotide to another pyrimidine (C:G ↔ T:A). Accordingly, when the *pyrF* sequence was analyzed in the few Ura^+^ mutants isolated from experiments with the controls strains, we found a significant enrichment of transversion mutations, e.g. 163(T→G), 164(A→T) and 164(A→C) (**Fig. 4c**). All these revertant (i.e. Ura^+^) clones had a very similar growth phenotype when grown in M9 minimal medium with glucose as the only carbon source, both when compared to each other or to their parental strain EM42 (**Fig. S3** in the Supporting Information). Interestingly, we could not isolate Ura^+^ mutants with the wild-type *pyrF* sequence (with a Lys residue at position 55 of PyrF; **Fig. 4b**) in any of these experiments. In agreement with our results, Long *et al*.^52^ showed that transition mutations are 16 to 82-fold more abundant than transversions in bacterial strains lacking a functional MMR system (both *Deinococcus radiodurans* and *P. fluorescens*), in sharp contrast to the mere < 3-fold found in the wild-type strains (i.e. spontaneously occurring). Horst *et al*.^24^ also indicated that DNA transitions and frameshift mutations were more abundant in *E. coli* cells lacking a functional MMR system. Regardless of the nature of the mutations introduced by these tools, these experiments show that the conditional mutator devices can be used to accelerate the emergence of different phenotypes. However, a major limitation of this set of plasmid-borne devices is the difficulty of curing them from the target cells, even in the absence of selective pressure. This shortcoming was fixed by constructing a new generation of ‘curable’ mutator devices as explained below.

### Design and validation of a new generation of plasmid-based, easy-to-cure mutator devices for Gram-negative bacteria

Previous attempts to cure isolated clones from the set of plasmids based on vectors pSEVA2311 and pSEVA2514 proved unsuccessful, even after >10 repeated passages of individual colonies under non-selective conditions (data not shown). This situation not only precludes precise temporal control of the accelerated evolution protocol, but also prevents the precise assessment of the (potential) occurrence of secondary mutations in the genome that do not have a selectable phenotype associated to their emergence. In particular, whole-genome sequencing needs to be performed to study the frequency and nature of mutations arising in conditional mutator strains, as well as the *global* mutation rates—as opposed to the *local* effects in individual genes that confer a macroscopic phenotype. Moreover, high-quality readings in whole-genome sequencing cannot be achieved if the cells carry plasmids (that would be co-purified with genomic DNA, and would interfere in the assembly process). In order to overcome this state of affairs, and due to the tedious work required to cure the mutator and control plasmids in all the strains previously tested, we decided to build a new version of easy-to-cure mutator systems using a technology recently developed in our laboratory. This methodology relies on the target curing of vectors by means of *in vivo* digestion mediated by the I-*Sce*I homing endonuclease^71^. For this purpose, we constructed vectors pS2311SG, pS2514SG, pS2311SGM and pS2514SGM by USER assembly (**Table 1** and **S1**). These standardized vectors, which are all derivatives of pSEVA2311, pSEVA2514, pS2311M and pS2514M, respectively, contain (i) an engineered I-*Sce*I recognition site that can be recognized and cleaved off by the endonuclease I-*Sce*I of *Saccharomyces cerevisiae*^72^ and (ii) a module for the constitutive expression of *msfGFP* (i.e. *P*_*EM7*_→*msfGFP*, where the gene encoding the monomeric superfolder GFP is placed under control of the synthetic *P*_*EM7*_ promoter) (**Fig. 5a**). This last module facilitates the selection of bacterial clones by examination of green-fluorescent colonies under blue light during the plasmid curation protocol. To this end, the accelerated mutagenesis protocol was upgraded by including a plasmid-curing step (**Fig. S4** in the Supporting Information). In this case, isolated clones are transformed with a helper plasmid that carries the gene encoding the I-*Sce*I endonuclease under control of an inducible expression system. Loss of the plasmid carrying the mutator device can be easily inspected as the corresponding colonies will also loss green fluorescence.

**Figure 5.**
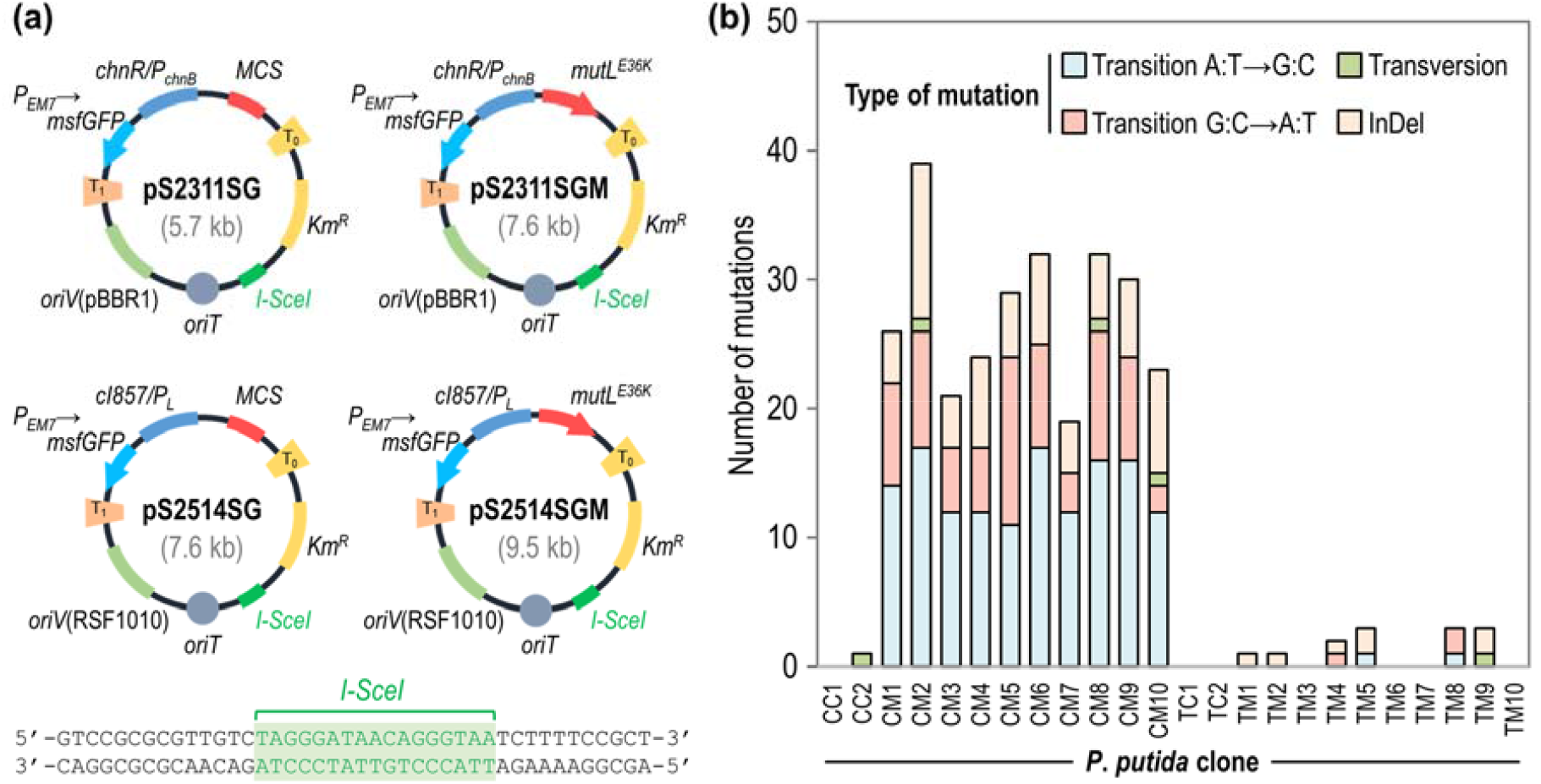
Design and implementation of a new generation of plasmid-based, easy-to-cure mutator devices for Gram-negative bacteria. (a) Plasmids pS2311SG, pS2514SG, pS2311SGM and pS2514SGM, derivatives of vectors pSEVA2311, pSEVA2514, pS2311M and pS2514M, respectively, were engineered with a I-SceI recognition site (indicated at the bottom of the figure) to render them compatible with the plasmid curing system based on targeted degradation mediated by the I-*Sce*I endonuclease^71^. These conditional mutator plasmids also carry a synthetic module for the constitutive expression of *msfGFP* (i.e. *P*_EM7_→*msfGFP*) that facilitate the selection of green fluorescent clones by examination of colonies under blue light. Functional elements in the plasmids not drawn to scale; *Km*^R^, kanamycin-resistance marker; *MCS*, standard multiple cloning site. **(b)** Mutation spectra caused by the mutator devices classified in functional categories. Control strains [i.e. *P. putida* KT2440/pS2514SG (TC) and KT2440/pS2311SG (CC)] and the conditional mutator strains [i.e. *P. putida* KT2440/pS2514SGM (CM) and KT2440/pS2311SGM (™)] were incubated in shaken-flasks in a non- selective culture medium. After 5 h (OD_600_ = 0.3), expression systems were induced thermally (in a water bath at 40°C for 15 min) or chemically (addition of 1 mM cyclohexanone). All cultures (induced and non-induced) were re-incubated at 30°C until reaching an OD_600_ = 0.6. Several dilutions of the cultures were plated on LB agar for isolation of individual colonies. After curing off the mutator and control (i.e. empty) plasmids from the respective clones, genomic DNA was isolated, sequenced and the readings assembled. The emergence of transition (A:T→G:C; G:C→A:T), transversion (A:T→T:A; G:C→T:A; A:T→C:G; G:C→C:G) and small insertion-deletion (InDel) events (< 50 bp) mutations was analyzed in the individual strains. Clones 1-5 and clones 6-10 were obtained from induced and non- induced cultures, respectively.

In these experiments, we firstly subjected the control strains (i.e. *P. putida* KT2440/pS2514SG and KT2440/pS2311SG) and the conditional mutator strains carrying the new set of plasmids (i.e. *P. putida* KT2440/pS2514SGM and KT2440/pS2311SGM) to the standard mutagenesis protocol to confirm the functionality of the easy-to-cure devices (**Fig. S4** in the Supporting Information). We investigated the emergence of Str^R^ mutants after implementing the accelerated mutagenesis protocol, and new induction conditions were tested to further characterize the tools. As expected, most of the recombinant strains harboring the easy-to-cure plasmids behaved quite similarly to the original strains carrying the first generation mutator devices (**Table S2** in the Supporting Information). The mutation frequency mediated by the mutator allele under control of the *c*I857/P_L_ expression system was essentially identical in all experiments, irrespective of whether the original or the upgraded set of plasmids was used. We detected a lower mutation frequency in strain KT2440/pS2311SGM (ca. 60% lower than the values observed in strain KT2440/pS2311M under similar experimental conditions). Such a trait was consistently accompanied by loss of green fluorescence in a significant proportion of the bacterial colonies isolated in solid medium, i.e. lysogeny broth (LB) agar, with or without Str supplementation. This result could be due to multiple factors, e.g. accumulation of loss-of-function mutations in the *msfGFP* gene stimulated by the same mutator device or unexpected decay or loss of the mutator plasmid in the absence of selection pressure (i.e. plasmid-borne kanamycin resistance).

To investigate the hypothesis above, we repeated the accelerated mutagenesis protocol with strain KT2440/pS2311SGM while maintaining kanamycin selection on the plates. We observed that, in the presence of the selection pressure borne by the mutator plasmid, all the bacterial colonies maintained green fluorescence and the overall mutagenesis frequencies were significantly higher than in all previous experiments (e.g. 270-fold higher than in the experiments with the same plasmid but omitting kanamycin; **Table S2**). Under these experimental conditions, the cyclohexanone-inducible mutator plasmids appear to exhibit leaky expression of the *mutL*^*E36K*^ allele, which led to similar mutagenesis frequencies in the absence or presence of inducer (**Table S2**). This observation helps explaining why, in the absence of selection pressure, some cells may reduce the copy number of the pS2311SGM plasmid to alleviate the mutagenic effects caused by (semi) constitutive expression of *mutL*^*E36K*^—or even force complete plasmid loss in some clones. At the bacterial population level, this phenomenon could further translate into an overall decrease of the *global* mutagenesis frequency. This behavior was not observed in strain KT2440/pS2514SGM, which appears to exhibit a lower—but more tightly-regulated— expression level of the mutator allele than the ChnR/P_*chnB*_ counterpart (**Table S2**). Actually, extending the thermal induction of the *c*I857/P_L_-based mutator devices from 15 to 30 min did not affect the *global* mutagenesis frequency. In either case, the genetic upgrading of the plasmid toolbox was meant to facilitate the easy curing of the mutator devices, and the results of these experiments are explained in the next section.

### Easy-to-cure mutator devices enable a tight control of the global mutagenesis and reveal a wide landscape of genome modifications upon accelerated evolution

We decided to sequence the whole genome of several colonies isolated in non-selective medium (i.e. LB agar, 2-5 colonies for each experimental condition) in order to assess the frequency and nature of mutations mediated by the mutator devices. To this end, green-fluorescent colonies were selected after treatment (**Fig. S4** in the Supporting Information) and transformed with the helper pQURE6.L plasmid^71^, a conditionally-replicating vector that requires supplementation of 3-methylbenzoate (3-*m*Bz) to the culture medium to ensure plasmid maintenance (**Fig. S5** in the Supporting Information). In particular, plasmid pQURE6.L carries a synthetic module for the 3-*m*Bz–inducible expression of the *I-SceI* endonuclease gene (i.e. XylS/*Pm*→*I-SceI*; **Table 1**) and a second module for the constitutive expression of *mRFP* (i.e. P_14g_→*mCherry*), which, together, facilitate quick curing of mutator plasmids by positive selection of red- fluorescent colonies (**Fig. S4** in the Supporting Information; see also *Methods* for details on the curing procedure). In all cases, the mutator devices could be easily cured upon introduction of plasmid pQURE6.L. Moreover, this helper plasmid could be typically cured during a simple overnight incubation of individual colonies in LB medium without 3-*m*Bz (data not shown), similarly to the observations reported by Volke *et al*.^71^

Multiple colonies were isolated from the accelerated mutagenesis experiments using the upgraded mutator toolbox and, upon curing all plasmids, genomic DNA was extracted and purified prior to next generation sequencing. Whole-genome sequencing of genomic DNA enabled a precise elucidation of the nature of mutations elicited by these devices. In general, whole-genome sequencing data confirmed our previous findings, as the emergence of transitions was a clear signature of clones carrying the *mutL*^*E36K*^ allele in different configurations (**Fig. 5b** and **Table S3** in the Supporting Information). These single-nucleotide polymorphisms were largely non-synonymous, and transversions were observed to be extremely rare (i.e. 1 transversion per genome in a just a few isolated clones, no different from the frequency of transversions in any of the control strains). Importantly, the mutator devices also promoted the emergence of small insertion-deletion mutations (InDel, mostly consisting of 1-2 bp; **Fig. 5b** and **Table S4** in the Supporting Information). Frameshift insertions were the most abundant type of InDels detected in the isolated clones. Taken together, and consistently with the results of experiments reported in the previous section, the detailed exploration of mutations elicited by the cyclohexanone- inducible mutator devices indicate that this system promotes a nearly-constitutive mutator phenotype. This feature, in turn, triggers a relatively high mutagenesis frequency over short induction periods— probably caused by the leakiness observed for this system under these conditions. Finally, the transcriptional output afforded by the thermoinducible mutator plasmid seemed to be tightly-regulated. Thus, this device could be applied to long evolutionary experiments that alternate cycles of non- induction and induction of DNA mutagenesis coupled to selection of target phenotypes.

## CONCLUSION

In this work, we have constructed two synthetic biology devices to control the mutation rate in *P. putida*—and, due to the nature of the vectors used for these constructs, other Gram-negative bacteria as well—in a precise spatiotemporal fashion. We have interfered with the functioning of the endogenous MMR machinery by transiently overexpressing the endogenous dominant negative *mutL*^*E36K*^ allele of *P. putida*, thereby increasing mutation frequencies in multiple strains of *P. putida* by 2- to 438-fold under the conditions tested herein. Following a ‘mutagenesis-followed-by selection’ approach, we have successfully evolved three separate phenotypes arising from monogenic traits, i.e. resistance to the antibiotics Str and Rif and uracil prototrophy. Within this approach, we have firstly increased the genetic diversity in the bacterial population by inducing the activity of the synthetic mutator devices and, subsequently, isolated mutants onto a selective solid medium. In these experiments, the expression of the mutator *mutL*^*E36K*^ allele was driven from two inducible modules, i.e. the thermoregulated *c*I857/P_L_ and the cyclohexanone-regulated ChnR/P_*chnB*_ expression systems, which have been previously tailored for heterologous gene expression in different Gram-negative bacterial species. We observed that the mutation frequencies achieved with the cyclohexanone-inducible mutator devices (i.e. vectors pS2311M and pS2311SGM, which represent the first and second generation of the tools constructed in this study) were significantly higher than those obtained with the thermoinducible mutator counterparts (i.e. vectors pS2514M and pS2514SGM) for most of the experimental conditions tested. In agreement with previous studies conducted with *E. coli* and related species, we have also observed a higher emergence of transition and frameshift (InDel) mutations in cells displaying a temporarily-tampered MMR system^24^.

Interestingly, the cyclohexanone-triggered mutator devices afforded a significant level of leaky expression of *mutL*^*E36K*^, which in turn promoted a nearly-constitutive mutator phenotype that lead to high mutagenesis rates. The mutation frequencies achieved with this system were, however, lower than those reported with constitutive mutator strains where the mutator phenotype was originated by modifications in components of the endogenous MMR system. For example, Kurusu *et al*.^73^ reported that the frequency of occurrence of Rif^R^ mutants in a *Δ)mutS* derivative of *P. putida* KT2440 was 1,000- fold higher than that in the wild-type strain. Since mutation rates must be precisely controlled to avoid extensive accumulation of deleterious mutations and to prevent genomic instability, the overexpression of mutator alleles should be driven from tightly-regulated expression systems (which is always challenging, irrespective of the bacterial host^74^) or during short periods of time. Thus, the easy-to-cure mutator plasmids developed in this study, which can be rapidly removed from isolated clones displaying the phenotype of interest, offer a clear advantage over conventional mutator strains—where the mutator phenotype is elicited by genomic (hence, essentially irreversible) modifications, as epitomized by the emergence of mutator phenotypes of *P. aeruginosa* in clinically-relevant setups^75-77^. In contrast with the results of the ChnR/P_*chnB*_-dependent module, the thermoinducible mutator devices allowed for a tightly-regulated expression of *mutL*^*E36K*^. This tool may be applied to long evolutionary experiments that involves alternating cycles of non-induction and induction of mutagenesis coupled to phenotype selection (e.g. growth-coupled approaches). By modifying the induction conditions and the number of induction cycles, a landscape of mutation rates could be achieved and adapted to the needs of each evolutionary experiment. The control of these parameters might be crucial for accelerating the evolution of complex phenotypes in industrial MCFs, since it has been previously shown that microbial adaptation to specific stresses is favored with certain mutation rates^78^. Due to its particular metabolic architecture, this would likely be the case for *P. putida* as well^79^.

From a more general perspective, it should be noted that the MutL/MutS protein complex of the MMR machinery appears to be well-conserved in most bacterial species^80-81^. For instance, the MutS protein from *P. putida* and the MutL protein from *P. aeruginosa* were shown to functionally complement Δ*mutS* and Δ*mutL* mutants of *E. coli* and *Bacillus subtilis*, respectively^73,82^. Therefore, the broad-host-range mutator devices developed herein are expected to be functional in other bacterial hosts as well. In addition to their application for the accelerated evolution of phenotypes that depend on multiple mutations across the bacterial genome, the use of these devices also revealed an important feature of the MMR system relevant for synthetic biology. A number of genome modification approaches rely on specifically interfering with the bacterial MMR system to enable strand invasion^51,53,74,83^. Besides the intended modifications (e.g. as encoded in mutagenic oligonucleotides), there are several secondary mutations that could occur due to overexpression of mutagenic alleles. The tight spatiotemporal manipulation of this trait, afforded by the plasmids reported in this study, could enable a more precise control of genome modifications by restricting the mutation landscape to the intended alterations.

## METHODS

### Bacterial strains and growth conditions

The bacterial strains used in this work are listed in **Table 1**. *E. coli* DH5α was used for cloning and plasmid maintenance. *E. coli* and *P. putida* strains were routinely grown in lysogeny broth (LB) medium (10 g L^−1^ tryptone, 5 g L^−1^ yeast extract and 5 g L^−1^ NaCl) at 37°C and 30°C, respectively, in an orbital shaker at 150 rpm. For mutagenesis experiments, *P. putida* was grown in M9 minimal medium supplemented with 0.3% (w/v) glucose as the sole carbon source as indicated in the text. Cyclohexanone was added at 1 mM to cultures for induction of *mutL*^*E36K*^ expression as necessary. When appropriate, antibiotics were also added at the following concentrations (μg mL^−1^): gentamicin (Gm) 10; kanamycin (Km), 50; streptomycin (Str), 100; and rifampicin (Rif), 50. Supplementation of 20 μg mL^−1^ uracil was implemented to support bacterial growth of uracil-auxotrophic strains. Bacterial growth was estimated by measuring the optical density at 630 nm (OD_630_).

### General DNA manipulations and sequencing

Molecular biology techniques were performed essentially as described in standard protocols^84^. Oligonucleotides were purchased from Integrated DNA Technologies (IDT; Leuven, Belgium) and their sequences are provided in **Table S1** in the Supporting Information. DNA amplification was performed on a C1000 Touch^™^ Thermal Cycler (Bio-Rad Corp., Hercules, CA, USA) using Phusion *U* Hot Start DNA Polymerase or Phusion Hot Start II DNA Polymerase from Thermo Fisher Scientific (Waltham, MA, USA). DNA fragments were purified with a NucleoSpin^™^ Gel and PCR Clean-up kit (Macherey-Nagel, Düren, Germany). Restriction enzymes and T4 DNA ligase were obtained from Thermo Fisher Scientific and were used according to the supplier’s specifications. USER assembly was performed essentially as described by Nour-Eldin *et al*.^85^ with the commercial USER enzyme from New England BioLabs (NEB, Ipswich, MA, USA). Plasmid DNA was prepared with a NucleoSpin^™^ Plasmid EasyPure kit (Macherey-Nagel). *E. coli* chemical competent cells were prepared using the *Mix & Go E. coli* Transformation Kit from Zymo Research (Irvine, CA, USA). DNA amplification from a single colony (i.e. colony PCR) was performed with One *Taq* 2× Master Mix (NEB). Electrocompetent *P. putida* cells were prepared by washing twice an overnight culture of *P. putida* with 300 mM sucrose^86^. All cloned inserts and DNA fragments were confirmed by DNA sequencing (Eurofins Genomics, Ebersberg, Germany).

### Construction of broad-host range mutator expression vectors

Plasmid pSEVA2514-*rec2- mutL*_*E36K*_^PP^, described by Aparicio *et al*.^54^, was double-digested with XbaI and HindIII to obtain a 1.9-kb DNA fragment corresponding to the dominant-negative mutator allele *mutL*^*E36K*^ of *P. putida* KT2440. The purified DNA fragment was subsequently ligated with the pSEVA2541 and pSEVA2311 vectors, previously digested with the same restriction enzymes, to generate plasmids pS2514M and pS2311M, respectively. The easy-to-cure plasmids pS2514SG, pS2514SGM, pS2311SG and pS2311SGM, were subsequently constructed by USER assembly with the primers indicated in **Table S1**. These vectors contain an engineered I-*SceI* recognition site and an *msfGFP* gene under the control of the constitutive P_EM7_ promoter (**Fig. S4** in the Supporting Information), that make them compatible with the plasmid curation approach recently developed by Volke *et al*.^71^.

### Accelerated evolution experiments with *P. putida* recombinant strains carrying mutator plasmids

Overnight pre-cultures of the conditional mutator strains [e.g. *P. putida* KT2440/pS2514(SG)M and KT2440/pS2311(SG)M], as well as of their respective control strains [e.g. *P. putida* KT2440/pS2514(SG) and KT2440/pS2311(SG)], were used to inoculate 25 mL of non-selective M9 minimal medium at an initial OD_600_ of 0.075. After 5 h of incubation in an orbital shaker at 30°C (OD_600_ = 0.3), the expression systems were induced thermally (by incubation at 40°C for 15 min in a water bath) or chemically (with addition of 1 mM cyclohexanone). The cultures were subsequently re- incubated at 30 °C with shaking and stopped after 1.5 h (early-exponential phase, OD_600_ = 0.6), 2 h (mid-exponential phase, OD_600_ = 1) or 24 h (stationary phase, OD_600_ = 3). Several aliquots of bacterial cultures were plated on selective solid medium (e.g. LB agar supplemented with 100 μg mL^−1^ Str or with 50 μg mL^−1^ Rif as appropriate) to determine the appearance of mutant cells (e.g. Rif^R^ or Str^R^) in the bacterial population. The total number of viable cells in the bacterial population was also estimated by plating dilutions of the cultures on non-selective medium (e.g. LB agar plates). After 32 h of incubation at 30 °C, the number of colony forming units (CFUs) in the different culture conditions was estimated by visual inspection of the plates (see **Fig. S2** in the Supporting Information for an example). At least two biological replicates and two technical replicates were performed for each bacterial strain and selective culture condition, respectively.

### Vector curing procedure for easy-to-cure plasmids carrying mutator devices

Overnight pre- cultures of green-fluorescent colonies isolated from evolution experiments were transformed by electroporation with plasmid pQURE6.L (**Table 1** and **Fig. S5**). Transformed cells were recovered in LB medium supplemented with 2 mM 3-*m*Bz during 2 h. Dilutions were then plated on LB agar supplemented with 10 μg mL^−1^ Gm and 1 mM 3-*m*Bz. Red-fluorescent colonies that had lost the mutator plasmids were easily isolated after 24-48 h of incubation at 30°C. For curing the helper pQURE6.L plasmid, overnight pre-cultures of red-fluorescent colonies were grown and dilutions were plated on non-selective medium (e.g. LB agar). Non-fluorescent colonies were selected after 24 h of incubation and were stored for further analysis. Loss of both plasmids in the selected colonies was further confirmed by Gm and Km sensitivity (**Fig. S4**).

### Genomic DNA purification, library construction, and whole genome sequencing (WGS)

DNA was purified using the PureLink^™^ Genomic DNA purification kit (Invitrogen, Waltham, MA, USA) from 2 mL of overnight LB cultures inoculated from cryostocks prepared after curing the plasmids from the strains. The genomic DNA of each sample was randomly sheared into short fragments of about 350 bp. The obtained DNA fragments were subjected to library construction using the *NEBNext*^™^ DNA Library Prep Kit (NEB), following the supplier’s specifications. Libraries quality control was performed with a Qubit® 2.0 fluorometer and an Agilent^™^ 2100 BioAnalyzer. Subsequent sequencing was performed using the Illumina NovaSeq^™^ 6000 PE150 platform. For quality-control purposes, paired reads with any one of the following characteristics were discarded: (i) read contains adapter contamination; (ii) uncertain nucleotides (*N*) constitute >10% of either read; (iii) low quality nucleotides (base quality less than 5, Q ≤ 5) constitute >50% of either read. Libraries construction, sequencing and subsequent data quality control was performed by Novogene Co. Ltd. (Cambridge, United Kingdom).

## SUPPORTING INFORMATION

**Table S1**. Oligonucleotides used in this work.

**Table S2**. Mutation frequencies estimated with the different versions of mutator plasmids created in this work.

**Table S3**. Distribution of single nucleotide polymorphisms (SNP) in evolved populations of *P. putida. Table S4*. Distribution of small insertion-deletion (InDel) mutations in evolved populations of *P. putida*.

**Figure S1**. Protein sequence alignment of the NH_2_-terminal region of MutL proteins from different bacteria.

**Figure S2**. Appearance of rifampicin- and streptomycin-resistant mutants in populations of *P. putida* KT2440 carrying a mutator device.

**Figure S3**. Growth profile of selected *P. putida* Ura^+^ mutants isolated in mutagenesis experiments.

**Figure S4**. Upgraded protocol for accelerated evolution of phenotypes using the new generation of easy-to-cure mutator devices.

**Figure S5**. Physical map of the helper pQURE6.L plasmid.

## AUTHOR CONTRIBUTIONS

L.F.C. and P.I.N. designed the experimental plan and the overall research project. L.F.C. and A.C. carried the experimental work and drafted the figures and the manuscript, with further contributions by P.I.N. All authors discussed the results and interpreted the experimental data.

## ACKNOWLEDGEMENTS

We thank V. de Lorenzo and his team (CNB-CSIC, Madrid, Spain) for sharing research materials and for enlightening discussions. The financial support from The Novo Nordisk Foundation (grants NNF10CC1016517 and *LiFe*, NNF18OC0034818), the Danish Council for Independent Research (*SWEET*, DFF-Research Project 8021-00039B), and the European Union’s Horizon 2020 Research and Innovation Programme under grant agreement No. 814418 (*SinFonia*) to P.I.N. is gratefully acknowledged. L.F.C. was supported by the European Union’s Horizon 2020 Research and Innovation Programme under the Marie Skłodowska-Curie grants agreements No. 713683 (*COFUNDfellowsDTU*) and No. 839839 (*DONNA*).

