## Supporting Information for "Spatiotemporal manipulation of the mismatch repair system of *Pseudomonas putida* accelerates phenotype emergence"

**Table S1.** **Oligonucleotides used in this work.**

| Oligonucleotide[a] | Sequence (5'→ 3') | Use |
| --- | --- | --- |
| $P_{EM7}$-msfGFP-UC-F | AGA CTG TUG ACA ATT AAT CAT CGG CAT AGT ATA TCG G | Construction plasmids pS2311SG, pS2514SG, pS2311SGM and pS2514SGM |
| $P_{EM7}$-msfGFP-UC-R | AAT TAA TTA TUT GTA GAG TTC ATC CAT GCC G | |
| BufferSeq-UC-F | AAC AGT CUA TAG AGG CAT CTA CTG CGT AGC | |
| BufferSeq-UC-R | AAA ATC TCT GAU TGT GTC TCA TTG CTC TC | |
| Backbone1-UC-F | ATC AGA GAT TTU CAA AAA ACA ATA GAG GAG ACT GAA TTT TC | |
| Backbone1-UC-R | ACC GCG GUC CAA TTA ATT ATT AGA AAA ATT C | |
| Backbone2-UC-F | ACC GCG GUC CGC GCG TTG TCT AGG GAT AAC AGG GTA ATC TTT TCC GCT GCA TAA CCC TGC | |
| Backbone2-UC-R | AAT AAT TAA TUA AAG GCA TCA AAT AAA ACG AAA GGC TC | |
| Backbone1-UC-F-2514 | ATC AGA GAT TTU CAG CCA AAC GTC TCT TCA GGC CAC | |
| PS1 | AGG GCG GCG GAT TTG TCC | Check pSEVA plasmids cargo |
| PS2 | GCG GCA ACC GAG CGT TC | |
| *pyrF*-seq-F | GCA GTT TCT TGG CCG TGT CGC | Sequencing of *pyrF* gene |
| *pyrF*-seq-R | CCT TCC CGG AGA CCT TGT TGA AG | |

[a] Oligonucleotides labeled as 'UC' were designed for USER cloning.

**Table S2.** **Mutation frequencies estimated with the different versions of mutator plasmids created in this work.**[a]

| Mutator device and condition | pS2311(SG) | | pS2311(SG)M | | pS2514(SG) | | pS2514(SG)M | | |
|---|---|---|---|---|---|---|---|---|---|
| | Cyclohexanone | | | | Thermal induction | | | | |
| | 0 mM | 1 mM | 0 mM | 1 mM | 0 min | 15 min | 0 min | 15 min | 30 min |
| Plasmid 1.0[b] | – | $1.3\times10^{-6}$ | $1.2\times10^{-5}$ | $1.2\times10^{-5}$ | – | $3.8\times10^{-7}$ | – | $5.4\times10^{-6}$ | – |
| Plasmid 2.0[c] | $1.2\times10^{-6}$ | – | $2.3\times10^{-6}$ | $3.9\times10^{-6}$ | $8.7\times10^{-7}$ | – | $8.9\times10^{-7}$ | $2.2\times10^{-6}$ | $2.4\times10^{-6}$ |
| Plasmid 2.0[d] | – | – | $2.4\times10^{-4}$ | $2.7\times10^{-4}$ | – | – | – | – | – |

[a] Control strains [i.e. *P. putida* KT2440/pSEVA2514, KT2440/pS2514SG, KT2440/pSEVA2311 and KT2440/pS2311SG] and the conditional-mutator strains [i.e. *P. putida* KT2440/pS2514M, *P. putida* KT2440/pS2514SGM, KT2440/pS2311M and KT2440/pS2311SGM] were incubated in shaken-flask cultures in non-selective media. After 5 h ($OD_{600}$ = 0.3), the expression systems were induced either thermally (incubation in a water bath at 40°C for 15 or 30 min) or chemically (addition of 1 mM cyclohexanone). All cultures (induced and non-induced) were re-incubated at 30°C until reaching an $OD_{600}$ = 0.6. Several aliquots of bacterial cultures were plated on a selective medium [i.e. LB agar supplemented with 100 μg $mL^{-1}$ streptomycin (Str)] to assess the emergence of Str resistant ($Str^R$) mutants upon evolution. The total number of viable cells was estimated by plating dilutions of the cultures onto LB agar. Absolute mutation frequencies (i.e. number of $Str^R$ mutant cells per viable cells) were determined based on these figures.

[b] *Plasmid 1.0* indicates the first generation of mutator devices, i.e. plasmids pSEVA2311, pS2311M, pSEVA2514 and pS2514M.

[c] *Plasmid 2.0* indicates the second generation of mutator devices, i.e. plasmids pS2311SG, pS2311SGM, pS2514SG and pS2514SGM.

[d] Cultures plated on LB agar medium with kanamycin (selection pressure for plasmid maintenance).

**Table S3.** **Distribution of single nucleotide polymorphisms (SNP) in evolved populations of *P. putida*.**[a]

| Mutations/ Strain | Upstream[b] | Exonic[c] | | | | Downstream[d] | Upstream/ Downstream[e] | Transition[f] | Transversion[g] | Total[h] |
|---|---|---|---|---|---|---|---|---|---|---|
| | | Stop gain | Stop loss | Synonymous | Non-synonymous | | | | | |
| **CC1** | 0 | 0 | 0 | 0 | 0 | 0 | 0 | 0 | 0 | 0 |
| **CC2** | 0 | 0 | 0 | 1 | 0 | 0 | 0 | 0 | 1 | 1 |
| **CM1** | 0 | 0 | 0 | 5 | 17 | 0 | 0 | 22 | 0 | 22 |
| **CM2** | 0 | 0 | 0 | 7 | 17 | 0 | 3 | 26 | 1 | 27 |
| **CM3** | 0 | 0 | 0 | 5 | 12 | 0 | 0 | 17 | 0 | 17 |
| **CM4** | 0 | 0 | 0 | 5 | 12 | 0 | 0 | 17 | 0 | 17 |
| **CM5** | 0 | 0 | 0 | 8 | 16 | 0 | 0 | 24 | 0 | 24 |
| **CM6** | 1 | 0 | 0 | 7 | 17 | 0 | 0 | 25 | 0 | 25 |
| **CM7** | 0 | 0 | 0 | 1 | 13 | 0 | 1 | 15 | 0 | 15 |
| **CM8** | 0 | 0 | 0 | 2 | 22 | 0 | 3 | 26 | 1 | 27 |
| **CM9** | 0 | 0 | 0 | 6 | 16 | 0 | 2 | 24 | 0 | 24 |
| **CM10** | 0 | 0 | 0 | 4 | 11 | 0 | 0 | 14 | 1 | 15 |
| **TC1** | 0 | 0 | 0 | 0 | 0 | 0 | 0 | 0 | 0 | 0 |
| **TC2** | 0 | 0 | 0 | 0 | 0 | 0 | 0 | 0 | 0 | 0 |
| **TM1** | 0 | 0 | 0 | 0 | 0 | 0 | 0 | 0 | 0 | 0 |
| **TM2** | 0 | 0 | 0 | 0 | 0 | 0 | 0 | 0 | 0 | 0 |
| **TM3** | 0 | 0 | 0 | 0 | 0 | 0 | 0 | 0 | 0 | 0 |
| **TM4** | 0 | 0 | 0 | 0 | 1 | 0 | 0 | 1 | 0 | 1 |
| **TM5** | 0 | 0 | 0 | 0 | 1 | 0 | 0 | 1 | 0 | 1 |
| **TM6** | 0 | 0 | 0 | 0 | 0 | 0 | 0 | 0 | 0 | 0 |
| **TM7** | 0 | 0 | 0 | 0 | 0 | 0 | 0 | 0 | 0 | 0 |
| **TM8** | 0 | 0 | 0 | 2 | 1 | 0 | 0 | 3 | 0 | 3 |
| **TM9** | 0 | 1 | 0 | 0 | 0 | 0 | 0 | 0 | 1 | 1 |
| **TM10** | 0 | 0 | 0 | 0 | 0 | 0 | 0 | 0 | 0 | 0 |

[a] Control strains [i.e. *P. putida* KT2440/pS2514SG (TC) and KT2440/pS2311SG (CC)] and conditional-mutator strains [i.e. *P. putida* KT2440/pS2514SGM (CM) and KT2440/pS2311SGM (TM)] were incubated in shaken-flask cultures in a non-selective medium. After 5 h ($OD_{600}$ = 0.3), the expression systems were induced either thermally (incubation in a water bath at 40°C for 15 min) or chemically (addition of 1 mM cyclohexanone) (i.e. induced conditions). All cultures (induced

and non-induced) were re-incubated at 30°C until reaching an $OD_{600}$ = 0.6. Several dilutions were plated on LB agar for isolating individual colonies. After curing the mutator and control plasmids in these clones, genomic DNA was isolated and fully sequenced. Clones 1-5 and clones 6-10 were obtained from induced and non-induced cultures, respectively. In this context, *SNP mutations* refer to the variation in a single nucleotide which may occur at some specific position in the genome, including transition and transversion of a single nucleotide.

[b] *Upstream*: SNPs located within 1 kb upstream from the transcription start site of the gene.

[c] *Exonic*: SNPs located in the exonic region; *Stop gain/loss*: a non-synonymous SNP that leads to the introduction/removal of a *STOP* codon at the variant site; *Synonymous*: single-nucleotide mutation without changing the amino acid sequence; *Non-synonymous*: single-nucleotide mutation with changing amino acid sequence.

[d] *Downstream*: SNPs located within 1 kb downstream from the transcription termination site.

[e] *Upstream/Downstream*: SNPs located within the < 2 kb intergenic region, which is in 1 kb downstream or upstream of the genes.

[f] *Transition*: Point mutation that changes a purine nucleotide to another purine (A:T ↔ G:C) or a pyrimidine nucleotide to another pyrimidine (C:G ↔ T:A).

[g] *Transversion*: Substitution of a (two-ring) purine for a (one-ring) pyrimidine or vice versa.

[h] *Total*: Total number of detected SNPs.

**Table S4.** **Distribution of small insertion-deletion (InDel) mutations in evolved populations of *P. putida*.**[a]

| Mutations/ Strain | Upstream[b] | Exonic[c] | | | | | | Downstream[d] | Upstream/ downstream[e] | Insertion | Deletion | Total[f] |
|---|---|---|---|---|---|---|---|---|---|---|---|---|
| | | Stop gain | Stop loss | Frameshift deletion | Frameshift insertion | Non-frameshift deletion | Non-frameshift insertion | | | | | |
| **CC1** | 0 | 0 | 0 | 0 | 0 | 0 | 0 | 0 | 0 | 0 | 0 | 0 |
| **CC2** | 0 | 0 | 0 | 0 | 0 | 0 | 0 | 0 | 0 | 0 | 0 | 0 |
| **CM1** | 1 | 0 | 0 | 0 | 1 | 0 | 0 | 1 | 1 | 3 | 1 | 4 |
| **CM2** | 0 | 1 | 0 | 1 | 7 | 0 | 0 | 1 | 2 | 10 | 2 | 12 |
| **CM3** | 0 | 0 | 0 | 0 | 2 | 0 | 0 | 1 | 1 | 4 | 0 | 4 |
| **CM4** | 0 | 0 | 0 | 0 | 5 | 0 | 0 | 2 | 0 | 7 | 0 | 7 |
| **CM5** | 0 | 0 | 0 | 0 | 3 | 0 | 0 | 1 | 1 | 5 | 0 | 5 |
| **CM6** | 0 | 0 | 0 | 0 | 5 | 0 | 0 | 0 | 2 | 6 | 1 | 7 |
| **CM7** | 0 | 0 | 0 | 0 | 3 | 0 | 0 | 0 | 1 | 4 | 0 | 4 |
| **CM8** | 0 | 0 | 0 | 0 | 3 | 0 | 0 | 1 | 1 | 5 | 0 | 5 |
| **CM9** | 0 | 0 | 0 | 2 | 2 | 0 | 0 | 1 | 1 | 4 | 2 | 6 |
| **CM10** | 1 | 0 | 0 | 1 | 5 | 0 | 0 | 1 | 0 | 7 | 1 | 8 |
| **TC1** | 0 | 0 | 0 | 0 | 0 | 0 | 0 | 0 | 0 | 0 | 0 | 0 |
| **TC2** | 0 | 0 | 0 | 0 | 0 | 0 | 0 | 0 | 0 | 0 | 0 | 0 |
| **TM1** | 0 | 0 | 0 | 0 | 0 | 1 | 0 | 0 | 0 | 0 | 1 | 1 |
| **TM2** | 0 | 0 | 0 | 0 | 0 | 0 | 0 | 0 | 1 | 0 | 1 | 1 |
| **TM3** | 0 | 0 | 0 | 0 | 0 | 0 | 0 | 0 | 0 | 0 | 0 | 0 |
| **TM4** | 0 | 0 | 0 | 0 | 0 | 0 | 1 | 0 | 0 | 1 | 0 | 1 |
| **TM5** | 0 | 0 | 0 | 0 | 1 | 1 | 0 | 0 | 0 | 1 | 1 | 2 |
| **TM6** | 0 | 0 | 0 | 0 | 0 | 0 | 0 | 0 | 0 | 0 | 0 | 0 |
| **TM7** | 0 | 0 | 0 | 0 | 0 | 0 | 0 | 0 | 0 | 0 | 0 | 0 |
| **TM8** | 0 | 0 | 0 | 0 | 0 | 0 | 0 | 0 | 0 | 0 | 0 | 0 |
| **TM9** | 0 | 0 | 0 | 0 | 0 | 0 | 1 | 0 | 1 | 1 | 1 | 2 |
| **TM10** | 0 | 0 | 0 | 0 | 0 | 0 | 0 | 0 | 0 | 0 | 0 | 0 |

[a] Control strains [i.e. *P. putida* KT2440/pS2514SG (TC) and KT2440/pS2311SG (CC)] and conditional-mutator strains [i.e. *P. putida* KT2440/pS2514SGM (CM) and KT2440/pS2311SGM (TM)] were incubated in shaken-flask cultures in a non-selective medium. After 5 h ($OD_{600}$ = 0.3), the expression systems were induced either thermally (incubation in a water bath at 40°C for 15 min) or chemically (addition of 1 mM cyclohexanone) (i.e. induced conditions). All cultures (induced and non-induced) were re-incubated at 30°C until reaching an $OD_{600}$ = 0.6. Several dilutions were plated on LB agar for isolating individual colonies. After curing

the mutator and control plasmids in these clones, genomic DNA was isolated and fully sequenced. Clones 1-5 and clones 6-10 were obtained from induced and non-induced cultures, respectively. In this context, *InDel mutations* refer to the insertion or deletion of ≤ 50 bp sequences in the DNA.

[b] *Upstream*: InDels located within 1 kb upstream from the transcription start site.

[c] *Exonic*: InDels located in the exonic region; *Stop gain/loss*: InDels that lead to the introduction/removal of a *STOP* codon at the variant site; *Frameshift deletion/insertion*: InDel mutations changing the open reading frame with deletion or insertion; *Non-frameshift deletion/insertion*: InDel mutations without changing the open reading frame with deletion or insertion sequences of 3 or multiple of 3 bases.

[d] *Downstream*: InDel located within 1 kb downstream from the transcription termination site.

[e] *Upstream/downstream*: InDel located within the < 2 kb intergenic region, within the 1 kb downstream or upstream of the genes.

[f] *Total*: Total number of detected InDels.

**Figure S1. Protein sequence alignment of the $NH_2$-terminal region of MutL proteins from different bacteria.**

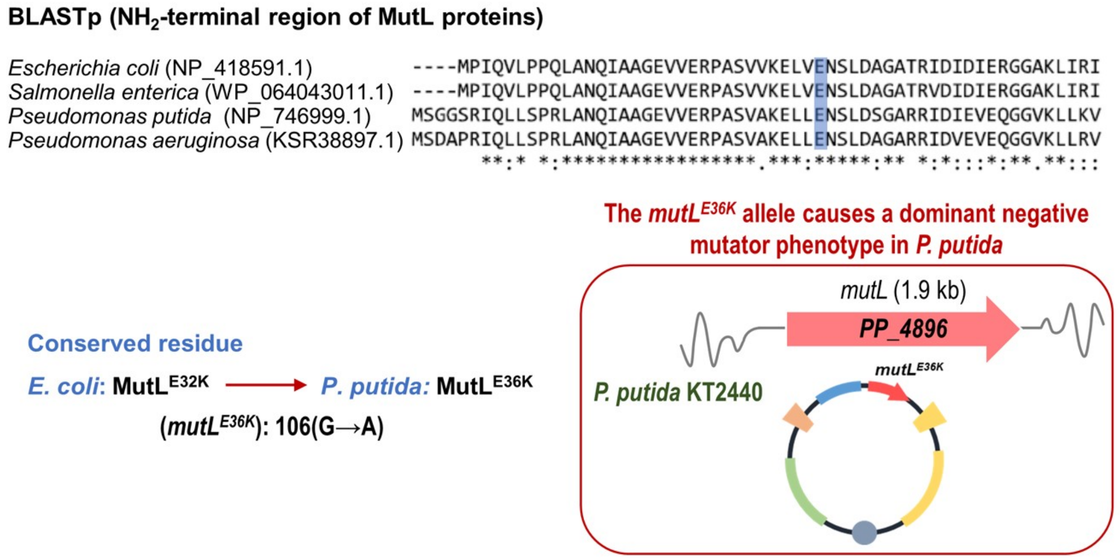


The overexpression of the dominant-negative mutator allele *mutL*$^{E32K}$ in *Escherichia coli* has been shown to increase the global mutation rate in this model bacterium (i.e. leading to a mutator phenotype)[1]. Based on protein sequence alignments of MutL proteins from several bacterial species, in this work we decided to utilize the *mutL*$^{E36K}$ allele from *P. putida* KT2440 for the manipulation of the mismatch repair system (MMR) of *Pseudomonas* sp.

**Figure S2.** **Appearance of rifampicin- and streptomycin-resistant mutants in populations of *P. putida* KT2440 carrying a mutator device.**

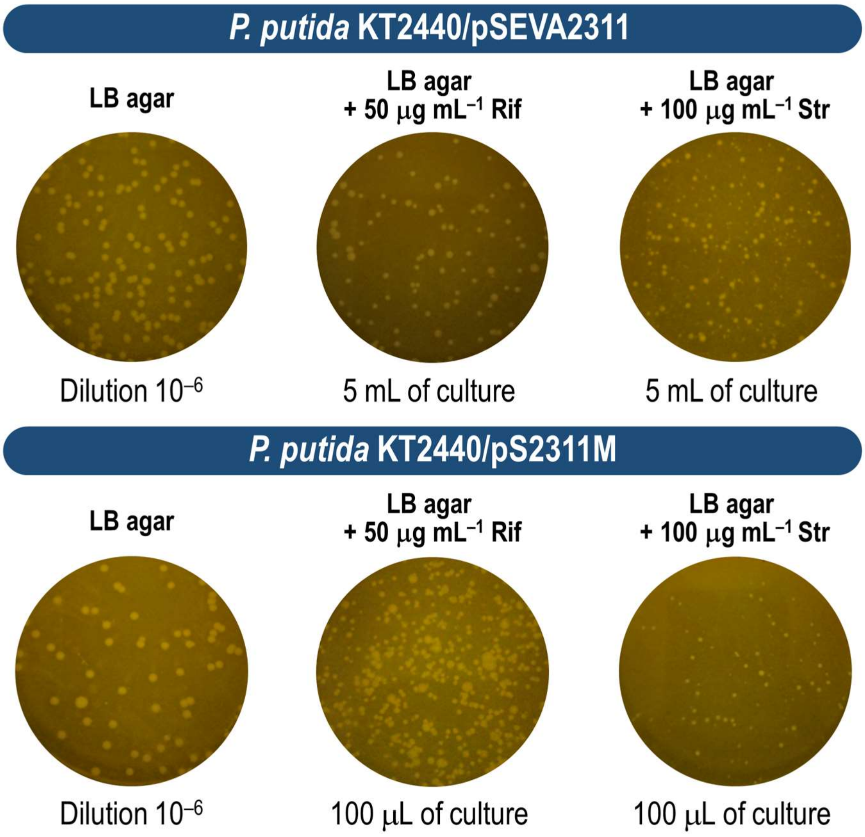


The expression of mutator devices in cultures of strains *P. putida* KT2440/pSEVA2311 (control, carries an empty vector) and KT2440/pS2311M was induced at an $OD_{600}$ = 0.3 by addition of 1 mM cyclohexanone. Cultures were harvested at an $OD_{600}$ = 0.6. Several aliquots and dilutions of the cultures were plated on selective medium [i.e. LB agar supplemented with rifampicin (Rif) or streptomycin (Str)] and non-selective medium (i.e. LB agar) as indicated in the figure and incubated at 30°C until individual colonies were clearly discernable. Representative Petri dishes from these experiments were photographed.

**Figure S3. Growth profile of selected *P. putida* Ura+ mutants isolated in mutagenesis experiments.**

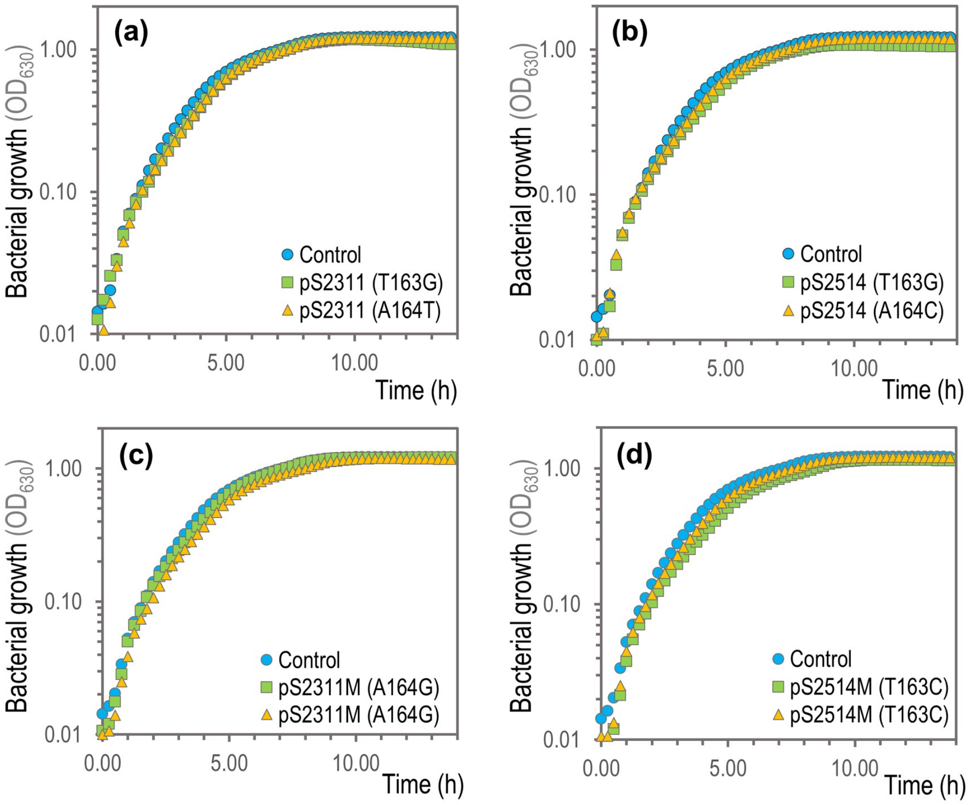


Bacterial growth of uracil prototrophic mutants (Ura+) derived from *P. putida* EM42 *pyrF* HM (Ura–) isolated from independent experiments where this strain was transformed with either the control vectors pSEVA2514 and pSEVA2311 **(a-b)** or the mutator devices (i.e. plasmids pS2514M and pS2311M) **(c-d)**. These experiments were conducted at 30°C in 96-well microtiter plates using M9 minimal medium containing 0.3% (w/v) glucose as the sole carbon source. Growth was estimated by measuring the absorbance at 630 nm ($OD_{630}$) every 15 min using a Synergy HI microplate reader (BioTek Instruments). *P. putida* EM42 was included in the growth experiments as a control (i.e. uracil prototrophic) strain. Mutations found in the *pyrF* gene (*PP_1815*) of the Ura+ mutants in these selected clones are indicated in parenthesis. Results represent the average values for $OD_{630}$ measurements from three biological replicates.

**Figure S4.** **Upgraded protocol for accelerated evolution of phenotypes using the new generation of easy-to-cure mutator devices.**

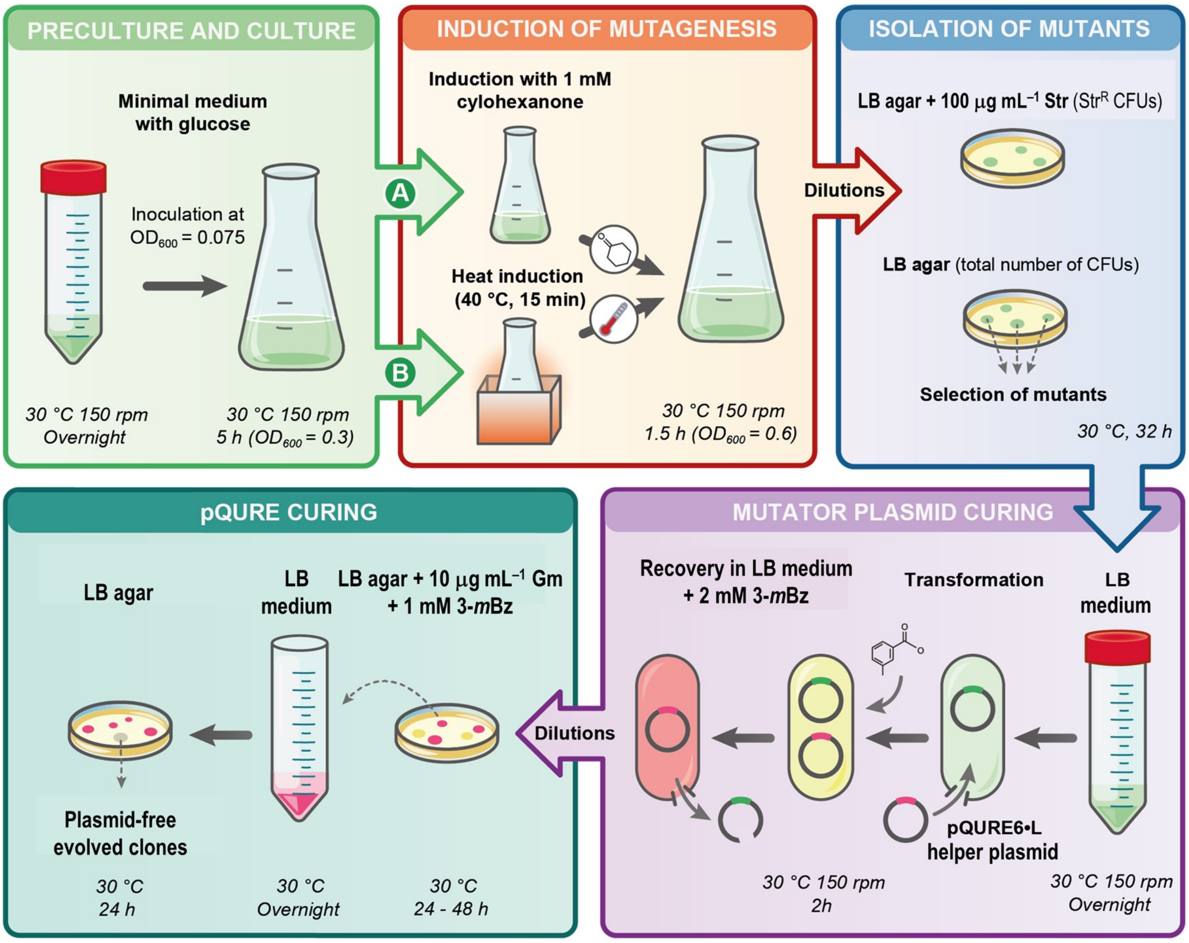


*P. putida* strains carrying the mutator devices inducible by cyclohexanone or temperature were evolved by following the upgraded mutagenesis protocol described in *Methods*. In short, aliquots or appropriate dilutions of the bacterial cultures were plated onto a selective medium (e.g. LB agar supplemented with 100 μg mL$^{-1}$ streptomycin) and non-selective medium (e.g. LB agar) to assess the appearance of mutants in the population and the total number of viable cells, respectively. All the colonies should present green fluorescence under the blue light due to the constitutive production of msfGFP, encoded in the second generation of mutator devices. Overnight pre-cultures of selected msfGFP$^{+}$ colonies were transformed by electroporation with the helper pQURE6·L plasmid, a conditionally-replicating vector dependent of the supplementation of 3-methylbenzoate (3-*m*Bz) to the culture medium to ensure its maintenance (see also **Fig. S5**). The helper plasmid carries, among other features, a synthetic module for the constitutive expression of *mCherry*, which facilitates the selection of red-fluorescent colonies by examination under blue light. Transformed cells were recovered on LB medium supplemented with 2 mM 3-*m*Bz during 2 h. Next, dilutions of the plasmid-bearing cultures were plated onto LB agar supplemented with 10 μg mL$^{-1}$ gentamicin (Gm) and 1 mM 3-*m*Bz. After 24-48 h of incubation at 30°C, red-fluorescent colonies that had lost the mutator plasmids (i.e. msfGFP$^{-}$) were easily selected by visual inspection of the plates. For curing the helper pQURE6·L plasmid, overnight pre-cultures of selected clones were prepared and plated on non-selective medium (e.g. LB agar). After 24 h, non-fluorescent colonies were isolated and stored for further analysis (e.g. whole-genome sequencing). Loss of both plasmids in the selected clones was further confirmed by Gm and kanamycin sensitivity.

**Figure S5.** **Physical map of the helper pQURE6·L plasmid.**

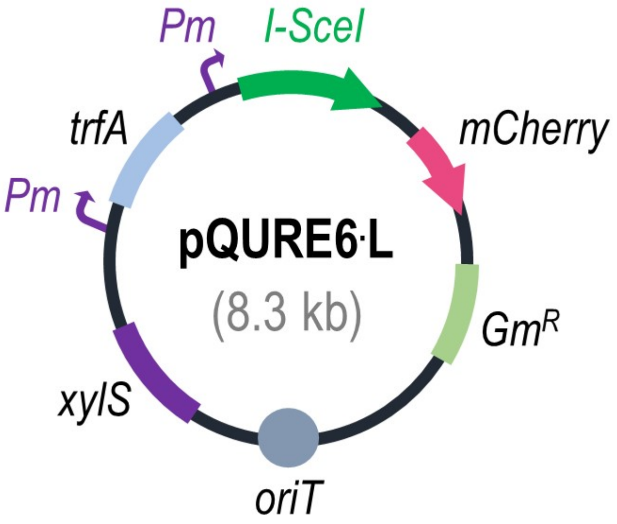


Plasmid pQURE6·L is a conditionally-replicating vector that requires the supplementation of the XylS/*Pm* inducer 3-methylbenzoate (3-*m*Bz) to the culture medium in order to be stably maintained by Gram-negative bacteria[2]. This plasmid bears a synthetic module for the 3-*m*Bz–inducible expression of the gene encoding the I-*Sce*I homing endonuclease (i.e. *Pm*→*I-SceI*) and the *trfA* gene of the low-copy-number *RK2* origin of replication (i.e. *Pm*→*trfA*), and a second module designed for constitutive expression of the red fluorescent protein mCherry (i.e. $P_{14g}$→*mCherry*). Together, these features facilitate quick curing of other (replicative) plasmids in the cell as long as they contain an I-*Sce*I recognition site. $Gm^R$, gentamicin-resistance marker.
